# Surface Tension and Stalk Elongation Drive *Dictyostelium* Morphogenesis

**DOI:** 10.64898/2026.09.02.748873

**Authors:** Seiya Nishikawa, Satoshi Kuwana, Gen Honda, Hidenori Hashimura, Satoshi Sawai, Shuji Ishihara

## Abstract

We investigate the mechanical principles underlying fruiting body morphogenesis in *Dictyostelium discoideum*. Quantitative shape analysis based on the Young–Laplace law, together with AFM indentation measurements, indicate surface tension as the dominant tissue-scale force acting on the culminating fruiting body. Based on this observation, we construct a hydrodynamic phase-field model with tunable surface and interfacial tensions, and analyze its behavior numerically. Our results show that, once a stalk begins to form, the elevation of the cell mass arises naturally through a dewetting process. Through quantitative comparisons with experimental measurements, we identify the mechanical conditions required for detachment from the substrate and for establishment of the characteristic morphology of the culminating fruiting body. Together, our model analysis highlights the importance of stalk-tip elongation and tissue-scale surface and interfacial tensions in the construction of large-scale three-dimensional tissues.

## I. INTRODUCTION

Multicellular organisms take on complex three-dimensional morphologies. The underlying tissues are, in many cases, multilayered and made up of heterogeneous cell types. Understanding how they form by coordinated cell-cell interactions requires not only the knowledge of the underlying biochemical signaling but also the mechanics [1–4]. A key question in morphogenesis is therefore how and where mechanical forces are generated and transmitted within tissues to give rise to desired changes in tissue shape. However, direct *in vivo* characterization aimed at uncovering how mechanical interactions shape three-dimensional structures remains limited to a handful of cases [5–9]. Given that complex morphological transformations can be decomposed into simpler modes of deformation, such as extension, bending, and branching, studying simple model systems should be effective in elucidating the relationship between mechanical forces and tissue morphology.

Fruiting body formation in the cellular slime mold *Dictyostelium discoideum* provides an excellent case for investigating the mechanics of three-dimensional tissue formation. Upon starvation, solitary cells cease dividing and aggregate by chemotaxis [10–13] and contact-dependent cell navigation [14, 15]. The resulting aggregate forms a mound within which cells begin to differentiate into two major cell-types: prestalk and prespore cells. The mound subsequently elongates to form a migratory slug with a distinct anterior-posterior polarity; prestalk cells are concentrated predominantly in the anterior region, whereas prespore cells occupy the posterior [16]. After a period of migration, the slugs transform into fruiting bodies through a process known as culmination, which involves vertical extension of the prestalk region, formation of the stalk and elevation of prespore cell mass (Fig. 1) [17, 18]. During the initial stage of culmination, anterior prestalk cells stop migrating whereas posterior prespore cells continue to move anteriorly. Concurrently, a subpopulation of posterior prestalk cells forms a basal structure called the basal disc. As the slug adopts a Mexican-hat-like morphology, and its prestalk–prespore axis become vertically reoriented, a subset of prestalk cells forms a central tip core and penetrate downward along the central axis to form a stalk. The stalk continues to elongate through prestalk-to-stalk differentiation at the anterior tip. The stalk is surrounded by cellulose-rich extracellular matrix that forms a rigid stalk tube. Differentiating stalk cells also accumulate cellulose on their surface and eventually become vacuolated. As the stalk elongates, the prespore region begins to detach from the substrate and the central basal disc. In the subsequent stage, the stalk continues to elongate while the prestalk and prespore cell masses are elevated. Although *Dictyostelium* belongs to the Amoebozoa, its development involves large-scale tissue deformation and several processes common to metazoan development, including cell differentiation, cell migration, and cell rearrangement [19, 20]. Nevertheless, the mechanical basis of cell mass detachment and elevation of the cell mass including the role of stalk tip elongation remains uncharacterized

**FIG. 1.**
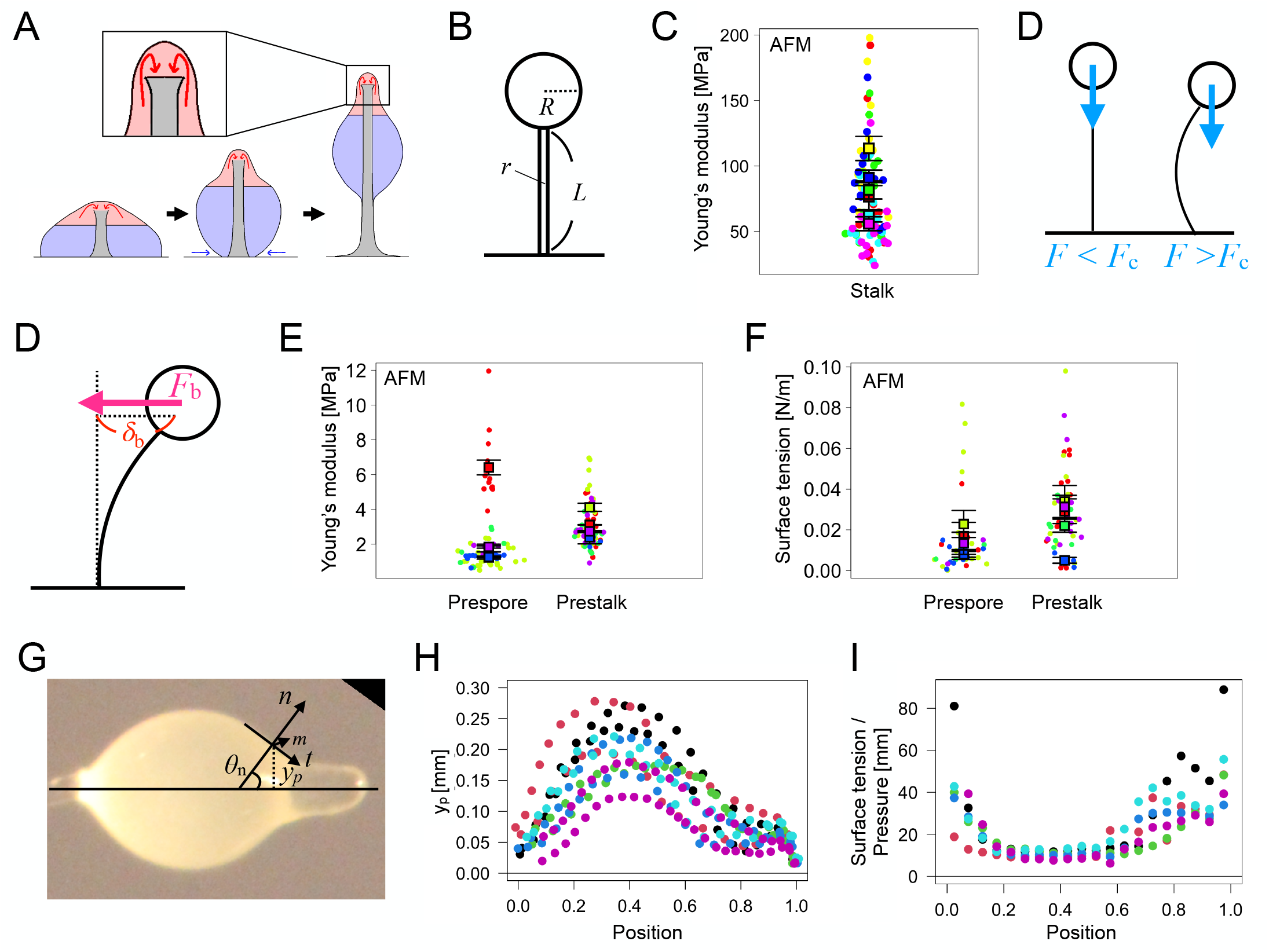
Quantification of forces acting on a *Dictyostelium* fruiting body. **A** A schematic diagram of the culmination process. Red, blue and gray masks indicate prestalk, prespore and the stalk region, respectively. Prestalk cells near the stalk tip enter the stalk from the top and differentiate into stalk cells (red arrow). **B** Geometric parameters *R, r*, and *L* denote the radius of the cell mass (approximated as a sphere), the radius of the stalk, and the stalk length from the substrate surface to the base of the cell mass, respectively. **C** AFM measurements of Young’s modulus of the stalk. Colors indicate individual samples (*n* = 6). Square markers and error bars are the mean and the standard error of the mean, respectively. **D** Stalk bending; the bending elastic force generated by the stalk *F*_b_ and the deflection *h*. **E, F** Estimated Young’s modulus (**E**) and the surface tension (**F**) of mature fruiting body. Colors, square markers, and error bars indicate individual samples (*n* = 3), the mean for each sample, and the standard error of the mean, respectively. Differences between prestalk and prespore were assessed by ANOVA (see SI Sec. S1 for details). **G** A representative snapshot of a fruiting body and the coordinate system used to calculate the surface tension. **t, n**, and **m** are unit vectors in the tissue-tangent direction, the tissue-normal direction, and the direction orthogonal to both **t** and **n**, respectively. *y*_p_ is the radial distance from the central axis to the surface. **H, I** *y*_p_ measured from experimental data (**H**) and the relative strength of surface tension calculated from the same images (**I**). Colors indicate individual samples (*n* = 6).

Fruiting body morphogenesis has previously been studied using mathematical modeling. Rubinow *et al*. developed a continuum model in which tissue-shape was determined by internal pressure and the boundary curvature [21]. Their model assumed that cells undergo chemotaxis in resonse to a cAMP gradient oriented along the prestalk–prespore axis. With appropriate chosen kinetics of cAMP production and degradation, the model reproduced the characteristic tissue contour observed during early culmination. Marée *et al*. subsequently developed a cellular Potts model (CPM) that describes differential cell-cell adhesion among multiple cell types [22, 23]. This model likewise assumed that cAMP signaling from the anterior tip and chemotaxis along the prestalk–prespore axis were the principal drivers of cell movement. It successfully reproduced the later stages of culmination, including elevation of the prespore mass and formation of a spore head atop a long, slender stalk. However, cell motility in the CPM was implemented through prescribed motility biases rather than explicit force-balanced interactions. Consequently, although the model reproduced the overall morphology, it did not enforce local force balance and could not determine the relative contributions of stalk-tip elongation and these biases to the elevation of the cell mass. Moreover, recent experimental findings challenge the assumed central role of cAMP signaling during the later stages of development. Although oscillatory cAMP signaling is indispensable during aggregation, its role in later stages remains debated, with some studies reporting that it gradually diminishes [24] and is no longer detectable in migrating slugs [25]. Furthermore, knockdown of the cAMP-synthesizing enzyme adenylyl cyclase A (ACA) does not prevent fruiting body formation [26]. Taken together, these findings indicate that both the mechanical basis of culmination and the role of cAMP signaling in this process remain poorly understood.

Recent studies in multicellular systems, both *in vitro* and *in vivo*, have highlighted the role of surface and interfacial tensions in organizing cellular assemblies over length scales of hundreds to thousands of micrometers [27–34], comparable to the size of a *Dictyostelium* fruiting body. These tensions represent the interfacial free-energy cost per unit area and act to minimize the corresponding interfacial area. At the cellular scale, the balance of interfacial tensions governs adhesion and wettability between cells and substrates, thereby determining whether cell collectives spread over a surface or retract from it [27, 28]. Mechanistically, such wetting behavior arises from cytoskeletal contractility, adhesion-mediated cell-cell interactions, and traction forces exerted on the substrate [27, 35]. These force-generating processes give rise to effective interfacial tensions and are collectively described as “active wetting” [28], linking tissue-scale dynamics to the underlying cellular mechanical processes. Although this theoretical framework was originally developed for *in vitro* culture systems, recent work has suggested that mesenchymal cell aggregation during mouse intestinal folding can be interpreted as a dewetting process in vivo [33]. In *Dictyostelium*, existing CPM models [22, 23, 36] implicitly encode interfacial tensions through contact energies, making it difficult to distinguish the contributions of individual interfaces to the overall morphogenetic movements. Notably, the cell mass of a mature fruiting body is nearly spherical, resembling a liquid droplet [18]. It has been shown that, when most of the prespore cells are replaced with mineral oil, the oil is still elevated while the overall shape of the cell mass is preserved [37]. These observations suggest that the large-scale morphology of the fruiting body may be governed primarily by the balance of tissue-scale interfacial tensions, rather than by the directed chemotactic migration of individual cells. However, the mechanical roles of the distinct interfaces in driving wetting, detachment, and elevation during culmination remain unclear.

Here, we develop a continuum mechanical framework in which tissue-scale interfacial tensions are formulated as explicit mechanical variables, allowing us to investigate how their balance governs wetting, detachment, and elevation during culmination. In Sec. II, we show that the tissue surface tension of the fruiting body is sufficiently large and that it plays a central role in shaping the characteristic morphology. We then construct a continuum phase-field model in Sec. III coupled to fluid dynamics while enforcing mechanical force balance. In Sec. IV, we compare simulations with experimental observations and show that the proposed mechanical framework explains elevation of the cell mass without requiring directed chemotactic migration. Finally, in Sec. V, we summarize our findings and discuss their biological implications.

## II. ESTIMATION OF FORCES ACTING ON A FRUITING BODY

### A. Gravity and mechanical stability of the stalk

To identify the dominant mechanical forces underlying fruiting body formation, we first measured the geometric and material properties of the tissue and estimated the magnitude of the relevant machanical forces. For simplicity, we approximated the shape of a fruiting body as a sphere with a central cylindrical beam as shown in Fig. 1B and assumed typical values for the stalk radius *r*, length *L*, and cell-mass radius *R*: 1.0 *×* 10^−5^ m, 7.1 *×* 10^−4^ m, and 1.0 *×* 10^−4^ m, respectively (Table S1). With the mass density of cells approximately 1.0 *×* 10^3^ kg/m^3^, the gravitational force acting on the sphere is approximately *F*_g_ = *ρ*_m_*V g* ≃ 4.1 *×* 10^−8^ N, with the sphere volume *V* ≃ 4.2 *×* 10^−12^ m^3^ and gravitational acceleration constant *g* = 9.8 m/s^2^.

Is the stalk sufficiently stiff to resist the gravitational loading? The stiffness of a beam is determined by the Young’s modulus *E* and the beam radius. Specifically, for a cylindrical beam with radius *r*, it is characterized by *EI* where *I* = *πr*^4^*/*4 represents the second moment of inertia of the cross-section[38]. We measured the Young’s modulus of the stalk by atomic force microscopy (AFM), and found *E* ≃ 8.0 *×* 10^7^ Pa (Fig. 1C; see Methods and Supplementary Information (SI) Sec. S1 for detailed protocols of the AFM analysis). For a beam with one end fixed, the critical Euler buckling force is *F*_c_ = *π*^2^*EI/*4*L*^2^ ≃ 3.1 *×* 10^−6^ N [39]. This is much larger than the gravitational force estimated above, indicating that the stalk is sufficiently stiff to avoid buckling. We next estimated bending stiffness of the stalk. When the free end of the stalk is subjected to a horizontal force *F*_b_ N, the end position shifts by a distance *δ*_b_ *µ*m, with the relationship *F*_b_ = (4*EI/L*^3^)*δ*_b_ ≃ 7.0 *×* 10^−9^*δ*_b_ N (Fig. 1D). Assuming a fruiting body that is placed horizontally in the air, the balance between the bending of the beam and gravitational forces on the sphere gives a deflection of only *δ*_b_ = 6.1 *×* 10^−6^ m, which is smaller than the stalk radius (10 *µ*m). These calculations indicate that gravity is negligible in fruiting body mechanics. Consistent with this notion, fruiting body formation is unaffected even when oriented horizontally or upside-down [40].

### B. Tissue surface tension

To assess the magnitude and regional variation of tissue surface tension, we performed two independent analyses. First, we estimated both tissue surface tension and Young’s modulus using AFM. For each fruiting body sample, we performed multiple indentations in the prespore and prestalk regions. By fitting the indentation curves to the empirical formula of [41], we obtained simultaneous estimates of Young’s modulus and surface tension (see SI Sec. S1; Fig. 1E,F). Young’s modulus was on the order of 10^0^ MPa, whereas surface tension was on the order of 10^−2^ N*/*m. Multiplying the measured surface tension by the characteristic perimeter of the fruiting body, *L*_sp_ = 2*πR* ≃ 6.2 *×* 10^−4^ m, the force arising from surface tension is estimated to be on the order of 10^−6^ to *×* 10^−5^ N. Thus, surface-tension forces exceed gravity (*F*_g_ ≃ 4.3 10^−8^ N) by more than one order of magnitude, and is comparable to or larger than the critical buckling force of the stalk (*F*_c_ ≃ 3.1 *×* 10^−6^ N). In addition, the measurements indicate that both Young’s modulus and surface tension are, on average, 1.5-2 times higher in the prestalk region than in the prespore region (Fig. 1E,F).

Because AFM-derived measurements of tissue-scale mechanics are known to depend on the size and the shape of the AFM probe [42], we complemented this analysis with an alternative approach based on tissue shape, which allowed us to evaluate surface tension relative to internal tissue pressure. In a cylindrically symmetric tissue, the surface tension *τ*_t_ and curvature *c*_m_ satisfy the Young–Laplace law, 2*τ*_t_*c*_m_ = *P*_0_, where *P*_0_ is the internal tissue pressure, *τ*_t_ denotes the surface tension along the tangential direction **t**, and *c*_m_ is the curvature in the bi-normal direction **m** [43]. Along the experimentally measured contour of a culminant (Fig. 1G), the curvature can be calculated by *c*_m_ = sin *θ*_n_*/y*_p_, where *θ*_n_ denotes the angle between the central axis and the local normal **n**, and *y*_p_ denotes the radial distance from the central axis to the surface. The relative magnitude of surface tension is thus given by

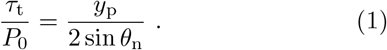

Figure 1H shows the radial distance measured from representative fruiting bodies. Fig. 1I shows the corresponding estimates of the relative tension, *τ*_t_*/P*_0_. Under the assumption of uniform internal pressure, the results indicate that the prestalk region has approximately two-fold higher surface tension than the prespore region. Note that in Fig. 1I, the relative tension also appears elevated at the rear end of the tissue, however, this is likely an artifact caused by the difficulty in distinguishing the boundary between the stalk and the prespore region in this area. Previous studies have reported internal cellular pressures in various cells, ranging from 10^1^–10^2^ N*/*m^2^ [44]. Assuming that tissue pressure of *Dictyostelium* is of the the same order as the internal pressure reported in other cell types yields an estimated tissue surface tension on the order of 10^−1^–10^0^ N*/*m approximately one order of magnitude larger than the AFM estimate.

Despite this quantitative difference, both independent approaches consistently indicate that the force generated by surface tension greatly exceed gravitational loading and comparable or greater than the stalk bending resistance. Notably, the estimated surface tension in *Dictyostelium* is two to four orders of magnitude higher than that typically reported for metazoan tissues (10^−5^ to 10^−2^ N*/*m) [45–51]. The unusually high surface tension may reflect the cellulose-rich extracellular matrix surrounding the culminant [52–55]. Furthermore, both approaches indicate that the surface tension in the pre-stalk region is approximately twice that in the prespore region. This regional contrast may arise from differences in tissue architecture, including the polarized, epithelial-like organization of cells at the culminant tip [56–58]. Overall, our analysis indicates that tissue surface tension is the dominant mechanical driver of tissue deformation during culmination.

## III. MATHEMATICAL MODEL FOR FRUITING BODY FORMATION

### A. Surface tension driven phase-field model

Motivated by the force estimates above, we formulate a continuum mechanical model to investigate how tissue surface tension, together with stalk-tip elongation, drives fruiting body morphogenesis. The governing equation satisfy mechanical force balance and are derived in detail in SI Sec. S2.

In our model, the prestalk–prespore cell mass, comprising prestalk cells in the anterior region and prespore cells in the posterior region, is represented as an incompressible viscous continuum, whereas the stalk along the central axis is represented as a prescribed rigid cylindrical boundary (Fig. 2A and B). The model consists of three variables as a function of time *t* and position ***r***: the phase field *ϕ*(*t*, ***r***), which describes the tissue domain and its interface; the tissue deformation velocity ***v***(*t*, ***r***); and the cell-type field *ρ*(*t*, ***r***), where *ρ* = 1 is prestalk and *ρ* = 0 is prespore (Fig. 2A). Here, *ϕ* ≈ 1 inside the tissue and *ϕ* ≈ 0 in the surrounding air, with the tissue boundary represented by a narrow interfacial region.

**FIG. 2.**
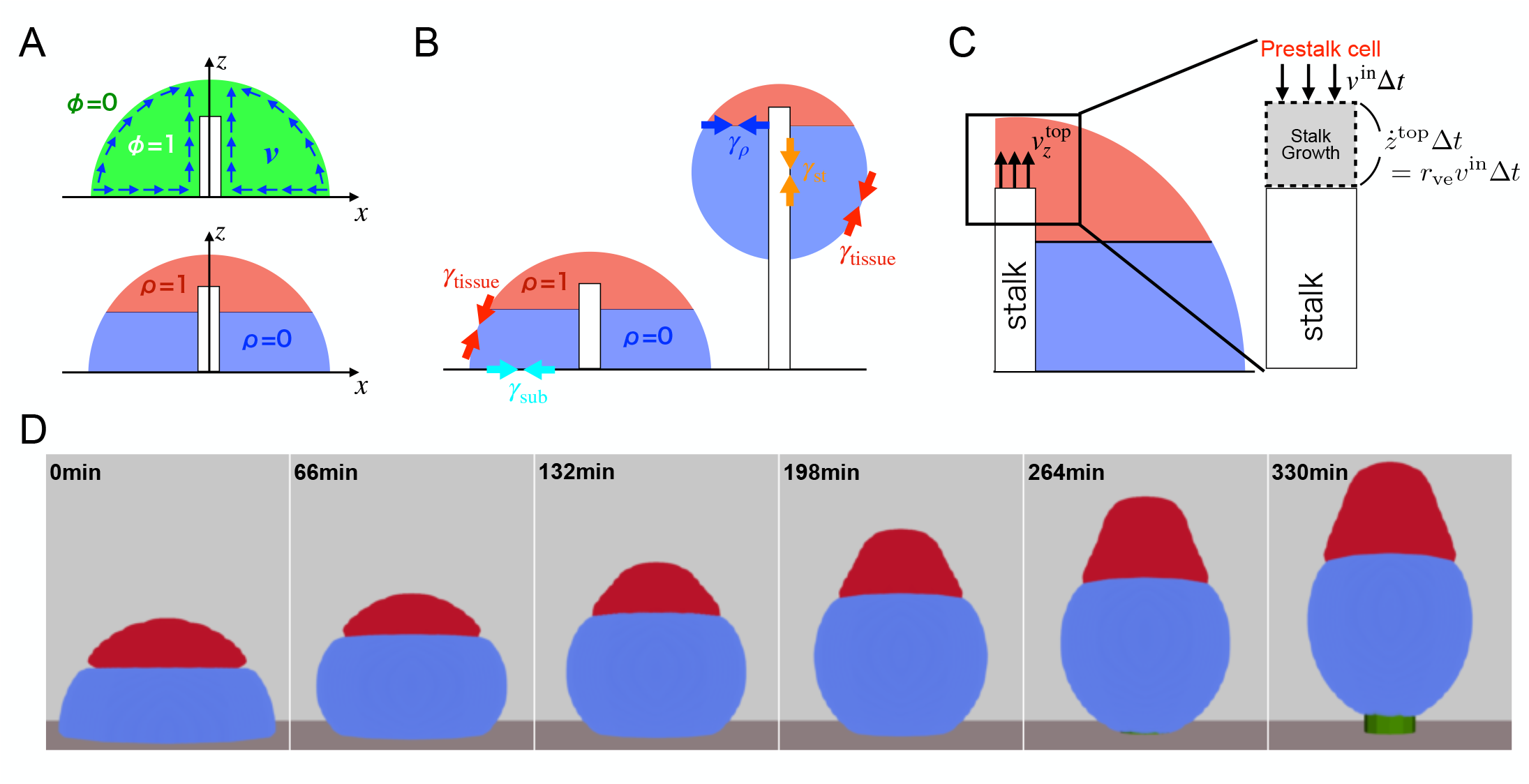
Numerical simulations of fruiting body formation. **A** Simulation framework. The phase field *ϕ* denotes the tissue interior (*ϕ* = 1) and exterior (*ϕ* = 0), respectively. *ρ* and 1 − *ρ* denote the fractions of prestalk and prespore cells, respectively. ***v*** denotes the tissue velocity field. **B** Surface and interfacial tensions in the model. *γ*_tissue_, *γ*_*ρ*_, *γ*_st_, and *γ*_sub_ denote the tissue surface tension, the prestalk–prespore interfacial tension, the tissue–stalk interfacial tension, and the tissue–substrate interfacial tension, respectively. **C** Boundary condition at the stalk anterior tip interface in the simulation. The stalk is represented as a vertical cylinder. Prestalk cells flow into the stalk tip at a constant speed *v*^in^, and upon differentiation the cell volume increases by a factor *r*_ve_, resulting in stalk growth. The parameters *r*_ve_ and *v*^in^ are fixed at 2.5 and 2 *×* 10^−5^, respectively, in all simulations. **D** Surface rendered images from a representative simulation (*γ*_pst_ = 5.7, *γ*_psp_ = 2.85, *γ*_*ρ*_ = 10.0, *γ*_sub_ = 1.0, and *γ*_st_ = 0.8, *η*_pst_*/η*_psp_ = 9.0).

Following the tension-corrected advected phase-field formulation [59], the time evolution of *ϕ* is given by

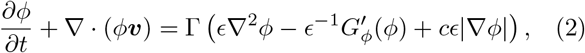

where 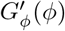 is the derivative of *G*_*ϕ*_(*ϕ*) = *ϕ*^2^(*ϕ* 1)^2^ with respect to *ϕ*, and *c* ≡ ∇ · ***n*** denotes the surface curvature, with ***n*** = −∇*ϕ/* |∇*ϕ*| being the outward unit vector normal to the tissue surface. The phase field separates into domains with *ϕ* = 0 and *ϕ* = 1, connected by an interface of width *O*(*ϵ*). The final term, *cϵ*|∇*ϕ*| , provides a curvature-dependent correction that cancels the leading-order spurious surface-tension contribution arising from the diffuse-interface regularization. This correction allows the tissue surface tension to be specified independently through the mechanical force balance introduced below, which determines the velocity field ***v*** that advects the tissue boundary.

The tissue domain represented by the phase-field *ϕ* = 1 is modeled as an incompressible viscous fluid with viscosity *η*_*v*_. The velocity ***v*** is governed by the Stokes equation as follows.

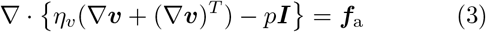

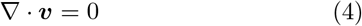

Here, *p*(*t*, ***r***) and ***I*** denote pressure field and identity matrix, respectively. ***f***_a_ represents the body-force force density acting on the tissue to be determined below. Eq. (4) represents the incompressibility of the tissue.

The third variable *ρ* is the cell-type index (prestalk *ρ* = 1 and prespore *ρ* = 0), so that *ρϕ* and (1 − *ρ*)*ϕ* represent the local volume fractions of prestalk and prespore cells at each position, respectively. Following the hydrodynamic formulation for two-component tissues [60], *ρϕ* obeys the continuity equation:

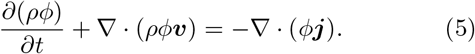

***j*** denotes the relative flux arising from differential motion between the prestalk and prespore cells. The prefactor *ϕ* confines this flux to the tissue domain, ensuring local conservation of both *ρϕ* and (1 − *ρ*)*ϕ*, except at the upper boundary of the stalk, where prestalk-cell transdifferentiation occurs (see below).

The model is formulated from a free-energy functional that accounts for phase separation between prestalk and prespore cells and for surface energy at the tissue, substrate and stalk interfaces:

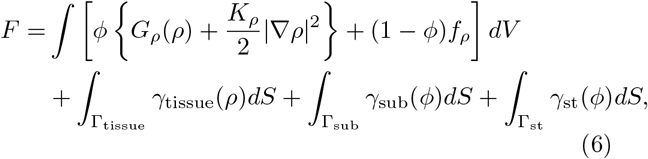

where *G*_*ρ*_(*ρ*) = *aρ*^2^(1 − *ρ*)^2^. The first term in Eq. (6) represents a volume integral of the free-energy density, with 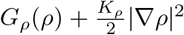 inside the tissue, and a fixed reference energy density *f*_*ρ*_ outside the tissue. The energy *G*_*ρ*_(*ρ*), with two local minima at *ρ* = 0 and *ρ* = 1, favors phase separation between prestalk <u>and p</u>respore cells. The interface has a thickness of 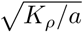 and a surface tension 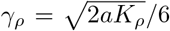. The second term in Eq. (6) represents surface energy along the boundary between the tissue and the air Γ_tissue_ (Fig. 2B). The surface integral over the entire interface can be written in the diffuse-interface form such that 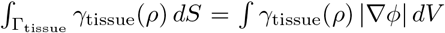. We set the tissue surface tension *γ*_tissue_(*ρ*) as

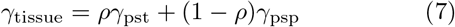

where *γ*_pst_ and *γ*_psp_ represent the surface tensions for the prestalk and prespore regions, respectively. The difference in surface tension contributes to maintaining the spatial segregation of prestalk and prespore regions. Similarly, the third and fourth terms represent the surface energies along the tissue–substrate interface Γ_sub_ and along the tissue–stalk interface Γ_st_, respectively, with *γ*_sub_(*ϕ*) and *γ*_st_(*ϕ*) denoting the corresponding surface tensions (Fig. 2B).

By requiring that the dynamics satisfy the second law of thermodynamics *dF/dt* ≤ 0, ***f***_a_ and ***j*** are obtained as

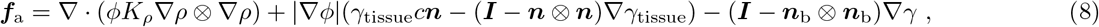

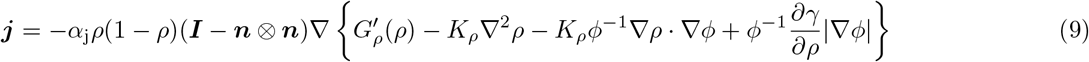

as described in SI Sec. S2.

The three terms in the force equation Eq. (8) represent prestalk-prespore interfacial tension, tissue surface tension and substrate/stalk interfacial tension, respectively. The first term in Eq. (8) is finite only on the surface between prestalk and prespore cells. In the second term, |∇*ϕ*| *γ*_tissue_*c****n*** represents the capillary force density localized at the tissue surface, directed along the surface normal ***n***, and proportional to the local surface curvature *c*, consistent with the Young–Laplace law. The remaining term represents the Marangoni force, arising from the spatial gradient of surface tension ∇*γ*_tissue_. In the third term, *γ* represents either *γ*_sub_(*ϕ*) or *γ*_st_(*ϕ*), depending on the position, while ***n***_b_ denotes the outward unit normal to the substrate boundary Γ_sub_ and the stalk surface Γ_st_. This term represents the tensile force acting at the corresponding interfaces. In Eq. (9), *α*_j_ denotes the mobility coefficient, and the quantity in braces is the chemical potential conjugate to *ρϕ, µ* = *ϕ*^−1^*δF/δρ*. The relative flux ***j*** is driven by the gradient of *µ*. The projection matrix ***I*** − ***n*** ⊗ ***n*** restricts diffusion of *ρ* to the local tangent plane of the tissue surface, thereby preventing normal flux across the tissue boundary.

Our model assumes that the stalk elongates as a result of an influx of trans-differentiating cells from the surrounding prestalk region at the tip region (the “AB core region” [61]). To represent this process, we defined a region at the upper end of the stalk (Fig. 2C; gray box) and considered the velocity component parallel to the stalk axis as 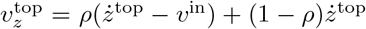. Here, *v*^in^ and *r*_ve_ are both constant and represent the influx of prestalk cells and a fixed dimensionless volumetric expansion ratio resulting from the cell-type transition, respectively. *ż*^top^ is stalk-tip velocity which is fixed at *ż*^top^ = *r*_ve_*v*^in^. We further assume *ϕ* = 1 and *ρ* = 1 on the upper-end of the stalk so that this region is always occupied by pre-stalk cells and remains in the cell mass. Accordingly, we imposed the following boundary conditions at the upper end of the stalk cylinder:

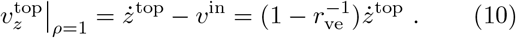

This boundary condition acts as a sink for prestalk cells. Although Eq. (5) indicates local conservation of the prestalk-cell density, *ρϕ*, within the tissue interior, the total prestalk volume, *ϕρ dV* , decreases as prestalk cells enter the stalk through the tip boundary and differentiate into stalk. On the lateral stalk wall and the substrate, we impose slip boundary conditions on the velocity field and zero-normal-gradient conditions on the scalar fields *ϕ* and *ρ*, namely, ∇*ϕ* · ***n***_b_ = 0 and ∇*ρ* · ***n***_b_ = 0.

In summary, we formulated a mathematical model that describes fruiting body morphogenesis. Equations (2)– (9) along with the boundary conditions described above, constitute a closed system that can be solved numerically. The time evolution of the shape variable *ϕ* is driven by surface and interfacial tensions as well as by the growth of the stalk tip and the subsequent reduction of the prestalk cell population. A key feature of the model is that mechanical equilibrium is enforced through the Stokes equation (Eq. (3)) , with surface and interfacial tensions incorporated as mechanical force densities acting within the tissue. The only active process in the model is the prescribed influx of prestalk cells at the tip, which drives stalk elongation. Moreover, the mechanical forces associated with surface and interfacial tensions are derived consistently from the free energy in Eq. (6) so as to satisfy the second law of thermodynamics.

### B. Simulation method

We numerically solved the three-dimensional model described above under the assumption of cylindrical symmetry. The governing variables were therefore represented by the axisymmetric fields *ϕ*(*t, r, z*), ***v***(*t, r, z*), and *ρ*(*t, r, z*) in cylindrical coordinates (*r, θ, z*). The computational domain was set to *L*_*r*_ = 50 and *L*_*z*_ = 150. The boundary conditions on the axisymmetric computational domain were specified as follows. At the symmetry axis, *r* = 0, we imposed *v*_*r*_ = 0 and ∂_*r*_*v*_*z*_ = ∂_*r*_*ϕ* = ∂_*r*_*ρ* = 0. At the outer radial boundary, *r* = *L*_*r*_, and the upper boundary, *z* = *L*_*z*_, zero-normal-gradient conditions were imposed for the scalar fields, ∇*ϕ* · ***n***_b_ = ∇*ρ* · ***n***_b_ = 0, where ***n***_b_ denotes the outward unit normal to the computational domain. Natural traction-free conditions, {*η*_*v*_(∇***v*** + (∇***v***)^*T*^) −*p****I***} ***n***_b_ = 0, were used for the velocity field.

Simulations were performed using the finite element method with the open-source PDE solver FreeFEM++ [62]. To ensure numerical stability, the phase-field equation, Eq. (2), was solved using the method proposed by Badillo [63, 64] (see SI Sec. S3 for details). The time step was set to Δ*t* = 0.1 simulation time unit. We employed the adaptive mesh refinement implemented in FreeFEM++ to adaptively refine the mesh every 100 simulation steps. The tissue shape was initialized as a hemisphere of radius 20, and the stalk radius *r*_st_ was fixed at 5 unless otherwise stated. The model parameters used in the present study are summarized in Supplementary Table S2. One simulation length unit corresponds to 9.0 *µ*m, and one simulation time unit corresponds to 0.3 s. The physical values of the surface and interfacial tensions were estimated by assigning the reference simulation value, 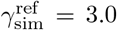 , to a representative tissue surface tension, 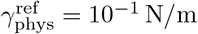. This gives 1 simulation tension unit ≃ 3.3 *×* 10^−2^ N*/*m. Note that this conversion provides an approximate measure of the physical scale rather than a precise calibration.

## IV. SIMULATION RESULTS AND COMPARISON WITH EXPERIMENTS

### A. Surface tension model reproduces tissue elevation

Figure 2D illustrates a representative numerical simulation (*γ*_pst_ = 5.7, *γ*_psp_ = 2.85, *γ*_*ρ*_ = 10.0, *γ*_sub_ = 1.0, and *γ*_st_ = 0.8). Starting from a hemispherical mound attached to the substrate, the basal area gradually shrinks while the prespore cell mass becomes increasingly rounded. As the stalk elongates, the prespore cell mass detaches from the substrate and is elevated (Movie S1). These results indicate that, within the present surface-tension-based framework, the prescribed prestalk-to-stalk transition at the tip region is sufficient to drive both the upward flow of prestalk and prespore cell mass and their detachment from the substrate. As shown below, successful tissue deformation and elevation depend critically on force balance at each of the three-phase contact lines.

### B. Pre-detachment stage analysis

To facilitate a detailed comparison between the simulation results and experimental observations, we divided the analysis of fruiting body morphogenesis into two stages: the pre-detachment stage (before tissue detachment from the substrate), and the elevation stage (during which the tissue ascends along the stalk). In the pre-detachment stage, we quantified tissue morphology by tracking the temporal evolution of three parameters: the contact angle *θ*, the basal contact radius *r*_bot_, and the tissue height *h* (Fig. 3A).

**FIG. 3.**
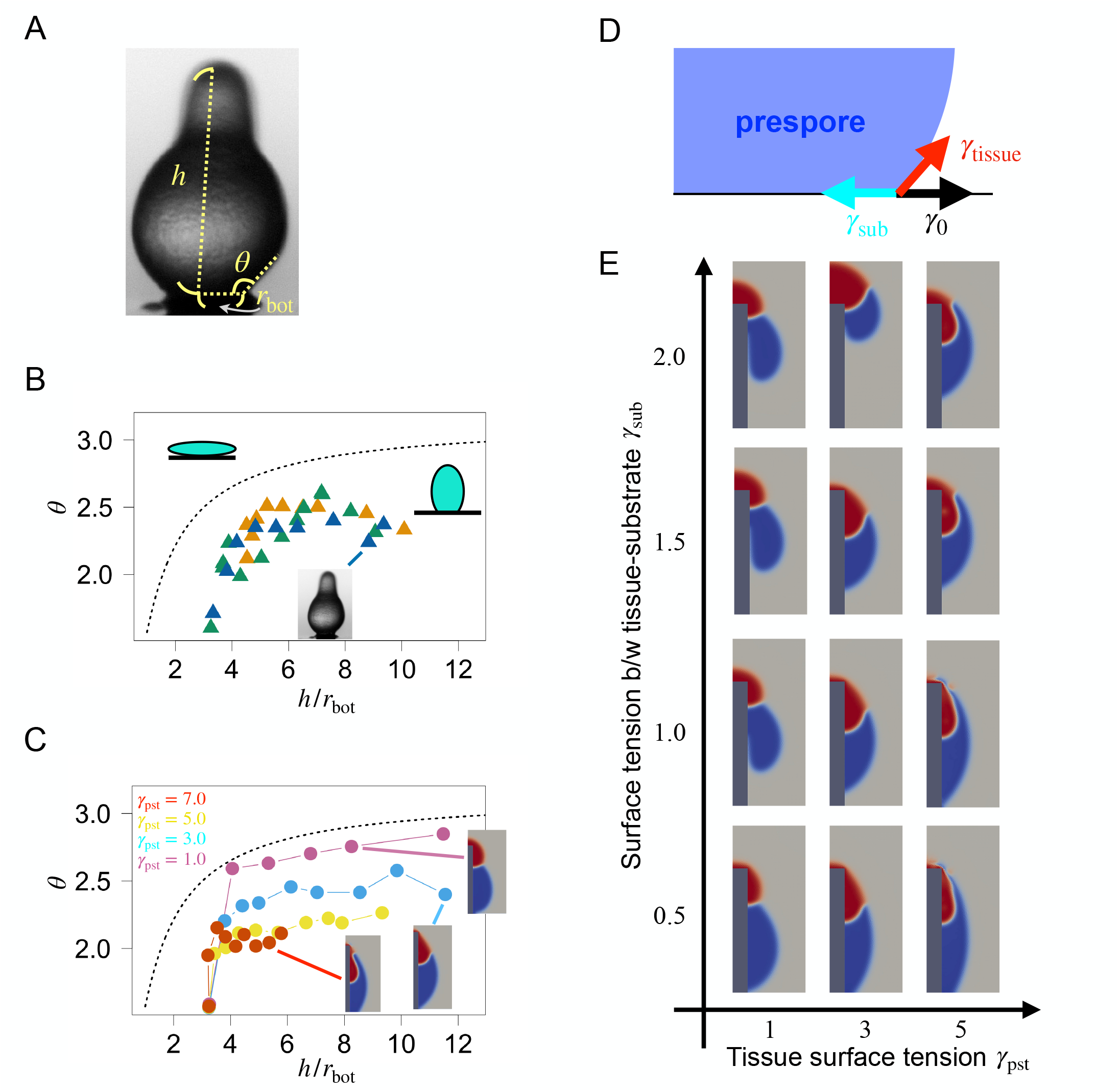
Dependence of tissue morphology and cell-mass detachment on tissue surface tension and tissue–substrate interfacial tension. **A** Geometric parameters: *r*_bot_, *θ*, and *h* represent the basal contact radius, the tissue contact angle, and the height from the substrate to the tip of the cell mass, respectively. **B, C** The ratio, *h/r*_bot_, versus contact angle *θ*; experimental data (B) and simulations (C). Developmental progression is from left to right. Colors indicate (B) individual samples (*n* = 3) and (C) the value of *γ*_pst_. The black dashed curves indicate the spherical-cap shape; tissues above the curves are more prolate, whereas those below are more oblate. For each simulation in **C**, the contact angle *θ* was manually measured every 50 simulation steps. **D** Schematic of the tension balance at the contact point. **E** Simulation results in the (*γ*_pst_, *γ*_sub_) plane at *t* = 250 min. Tissue shapes are shown as cross-sectional views in the *x*–*z* plane.

Figure 3B displays the experimental trajectories in the (*h/r*_bot_, *θ*) plane with different colors indicating three independent samples. As a geometric reference, we considered a spherical cap with a constant volume. The dimensionless relationship between the contact angle, the basal contact radius and the height of a spherical cap is given by

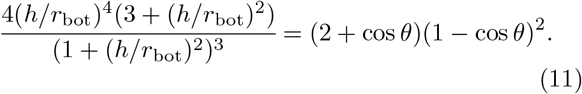

Shapes lying above the (*h/r*_bot_)-*θ* curve (Figs. 3B and C; black dashed curve) are more oblate, whereas those below are more prolate. After the initial rise in both the ratio *h/r*_bot_ and the contact angle *θ*, the increase in *θ* gradually slows down and appears to saturate, while *h/r*_bot_ continues to increase (Fig. 3B). Notably, the contact angle is smaller than that of a spherical cap with the same *h/r*_bot_, indicating that the tissue remains more vertically elongated than a spherical cap. The basal contact radius decreases until the cell mass detaches from the substrate, while the contact angle remains approximately constant.

We performed numerical simulations in which the initial tissue geometry was given as a half-ellipsoid with an experimentally estimated *h/r*_bot_. We varied the effective tissue surface tension, *γ*_tissue_ = *ργ*_pst_ + (1 − *ρ*)*γ*_psp_, while fixing the surface-tension ratio at *γ*_pst_*/γ*_psp_ = 2.0, consistent with the AFM measurements and tissue shape analysis described in Sec. II B. All other parameters were fixed at the values listed in Table S2. As shown in Fig. 3C, the simulated (*h/r*_bot_)-*θ* curves exhibited trends similar to those observed experimentally. The asymptotic contact angle varied with the effective tissue surface tension *γ*_tissue_.

For large *γ*_tissue_ (e.g., *γ*_pst_ = 7.0 in Fig. 3C red), *θ* takes a smaller value and does not reach the experimentally observed level. Conversely, when *γ*_tissue_ is small (e.g., *γ*_pst_ = 1.0, Fig. 3C magenta), the contact angle increases to a value larger than that observed experimentally. These results suggest that, in the early stage of fruiting body formation, *γ*_tissue_ takes an intermediate value. This point is discussed further in Sec. IV C.

### C. Detachment of the cell mass from the substrate

In the simulations, detachment of the cell mass from the substrate fails under certain parameter conditions. This observation is particularly interesting because there are mutants known to exhibit similar defects [65–69]. We thus examined how the tissue surface tension *γ*_tissue_, and the substrate-basal interfacial tension *γ*_sub_, with the air– substrate interfacial tension *γ*_0_ held fixed, determine tissue wettability and thereby control detachment dynamics.

Since an increase in *γ*_sub_ promotes shrinkage of the basal contact area (i.e., a decrease in *r*_bot_), while an increase in *γ*_tissue_ enhances resistance to tissue deformation, whether detachment occurs is determined by the balance among the interfacial tensions *γ*_sub_, *γ*_tissue_, and *γ*_0_ (Fig. 3D). Figure 3E summarizes the simulated tissue morphologies obtained from hemispherical initial conditions. Here, *γ*_pst_ and *γ*_sub_ were varied, while others were fixed (*γ*_pst_*/γ*_psp_ = 2.0, *γ*_*ρ*_ = 10.0, and *γ*_st_ = 0.8; see Table S2). For small *γ*_sub_, shrinkage of the basal contact area is limited and detachment does not occur. As *γ*_pst_ increases, the threshold value of *γ*_sub_ required for detachment increases. In the high-*γ*_pst_ regime, the prespore cells move upward around the outer surface of the prestalk region from below, progressively enveloping the prestalk cells. This occurs in both detaching and non-detaching cases; in the latter, excessive vertical elongation can cause numerical instability. For large *γ*_sub_, the basal surface contracts rapidly, leading to tissue detachment followed by elevation of the cell mass along the stalk. However, excessively large *γ*_sub_ leads to inward movement of prespore cells during the subsequent upward motion after detachment. The simulated morphology most closely resembled that of the experimentally observed morphology at intermediate values of *γ*_pst_ and *γ*_sub_, e.g., *γ*_pst_ = 3.0 and *γ*_sub_ = 1 (Fig. 3E).

The parameter sweep therefore indicates that detachment of the cell mass and its subsequent elevation require an appropriate balance between the tissue surface tension, varied through *γ*_pst_, and the tissue–substrate interfacial tension, *γ*_sub_. In the simulations described above, we fixed the prestalk-to-prespore surface-tension ratio *γ*_pst_*/γ*_psp_ = 2. Because no direct measurements of this ratio are available, we also examined a nearly uniform surface-tension condition (*γ*_psp_ = 0.95 *γ*_pst_ ; Fig. S2). The qualitative dependence of tissue detachment on *γ*_pst_ and *γ*_sub_ remained unchanged, indicating that the predicted detachment behavior is robust to uncertainty in the assumed surface-tension ratio.

### D. Elevation-stage morphological analysis

We next investigated the mechanical conditions required to reproduce the characteristic tissue morphology during the elevation stage. Necking of the prestalk region (Fig. 1G) is a prominent feature of fruiting-body morphology. Despite incorporating heterogeneous tissue surface tensions by setting *γ*_pst_*/γ*_psp_ = 2.0, the simulation shown above (Fig. 3E) exhibited less pronounced necking. This discrepancy indicates that the surface-tension difference alone is insufficient to account for the pronounced necking of the prestalk region. Previous studies have shown that prestalk cells form an epithelial-like columnar layer, whereas prespore cells lack such organization [56–58]. These observations suggest that the prestalk region has a lower fluidity and greater resistance to cell rearrangement than the prespore region. We therefore modeled the epithelial-like mechanical properties of the prestalk region by incorporating both the experimentally estimated surface-tension difference and a cell-type-dependent viscosity that accounts for lower fluidity and greater resistance to deformation and cell rearrangement. To represent this effect, we replaced the viscosity *η*_*v*_ in Eq. (3) with the cell-type-dependent viscosity *η*_*ρ*_(*ϕ, ρ*), defined as

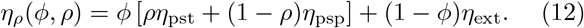

Here, *η*_pst_ and *η*_psp_ denote viscosities of the prestalk and prespore regions, respectively. To ensure computational stability, the viscosity of the exterior region *η*_ext_ was set equal to *η*_psp_. In these simulations, we varied the prestalk viscosity *η*_pst_ and prestalk surface tension *γ*_pst_ while keeping the prespore viscosity *η*_psp_ and prespore surface tension *γ*_psp_ fixed. Figure 4A summarizes the simulation results. Necking in the prestalk region, indicated by the black arrow in Fig. 4B, became more pronounced when the viscosity ratio *η*_pst_*/η*_psp_ increased. By comparison, varying *γ*_pst_*/γ*_psp_ had little effect on the overall tissue morphology and only subtly alters the neck shape within the explored parameter range.

**FIG. 4.**
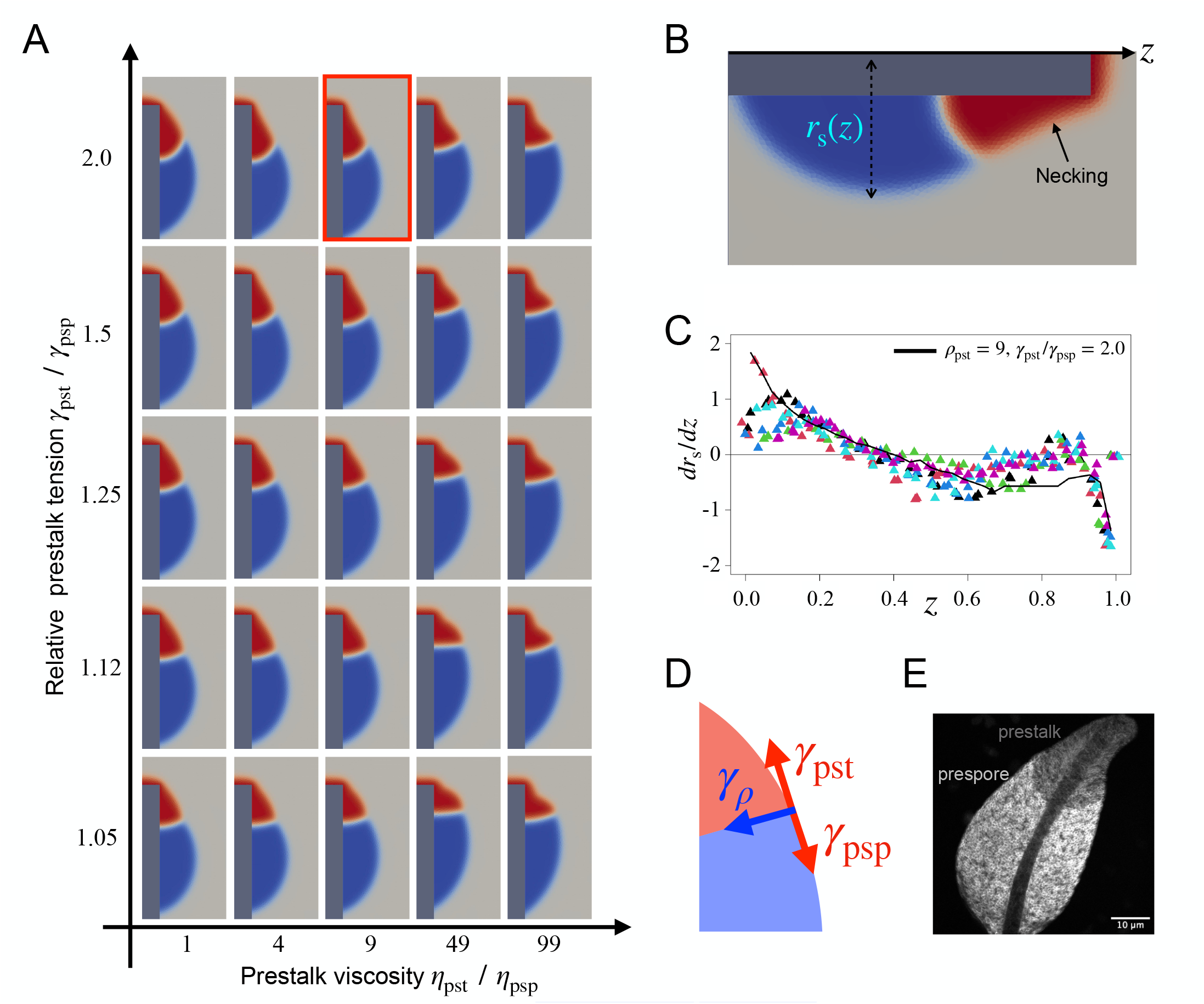
Tissue morphology depends on tissue viscosities and surface tensions. **A** Simulation results in the (*η*_pst_*/η*_psp_)-(*γ*_pst_*/γ*_psp_) plane at *t* = 250 min. The snapshot enclosed by the red box is taken from the same simulation as that shown in Fig. 2D. **B** Definitions of necking and the radius *r*_s_(*z*) used to quantify tissue shape. **C** Plot of normalized *z* versus *dr*_s_*/dz*. Points show experimental measurements, with colors indicating individual samples. The same experimental data as those shown in Fig. 1H were used. The black line shows the simulation result for the parameter set enclosed by the red box in **A**, which provides the best fit to the experimental data. **D** Schematic of the tension balance among *γ*_pst_, *γ*_psp_, and *γ*_*ρ*_. **E** Section image of a mid-culminant composed of plasma-membrane-labeled cells (PKBR1(N150)-mScarlet-I; see Methods for details), in which the brightly fluorescent and dim regions correspond to the prespore and prestalk domains, respectively.

For quantitative comparison, we measured the radial coordinate of the tissue boundary *r*_s_(*z*) as a function of *z*,, as shown in Fig. 4B, and calculated its derivative, *dr*_s_*/dz*, for each simulation and for the experimental data. In Fig. 4C, triangular markers denote the derivatives obtained from the experimental data shown in Fig. 1H, with different colors representing individual samples (*n* = 6). All experimental samples exhibited consistent morphological trends, including a rounded basal region and a tapered apical region with a distinct neck, indicating that our analysis captures a key feature of tissue shape. Among the simulated parameter sets (Fig. 4A), the combination *η*_pst_*/η*_psp_ = 9.0 and *γ*_pst_*/γ*_psp_ = 2.0 most closely reproduced the experimental boundary profile (black line in Fig. 4C), as quantified by the least-squares error between the simulated and measured *dr*_s_*/dz* values. The finding that the best-fitting parameter set features high viscosity and high surface tension in the prestalk region, supports our hypothesis that the prestalk tissue is mechanically more resistant to deformation and cell rearrangement, possibly reflecting its epithelial-like organization.

The influence of the surface tension ratio *γ*_pst_*/γ*_psp_ is most pronounced at the prestalk–prespore inteface, whose morphology is determined primarily by the balance among *γ*_pst_, *γ*_psp_, and *γ*_*ρ*_, (Fig. 4D). In the simulations shown in Fig. 4A, where *γ*_*ρ*_ is fixed, the prestalk– prespore interface became increasingly tilted and curved as *γ*_pst_*/γ*_psp_ increases, whereas a higher viscosity ratio *η*_pst_*/η*_psp_ tends to straighten it. Under the best-fitting parameter set (*η*_pst_*/η*_psp_ = 9.0 and *γ*_pst_*/γ*_psp_ = 2.0), the resulting prestalk–prespore interface was distinctly tilted and curved. This characteristic interfacial shape was also evident in experimental observations (Fig. 4E) [70], further indicating that our simulations capture the essential mechanical drivers of fruiting body morphogenesis.

### E. Influence of other interfacial tensions on tissue morphology

In this subsection, we numerically investigated how the remaining two interfacial tensions in our model, *γ*_*ρ*_ and *γ*_st_, influence tissue morphology during fruiting body formation. We performed two sets of simulations in which *γ*_pst_ was varied together with either *γ*_*ρ*_ or *γ*_st_ (Figs. 5A and B), while maintaining a fixed ratio *γ*_pst_*/γ*_psp_ = 2. As shown in these panels, large values of *γ*_st_ induce intrusion of the prespore region into the tissue interior, whereas excessively large *γ*_pst_ leads to incomplete detachment of the tissue from the substrate, consistent with the pre-detachment stage analysis in Sec. IV C.

**FIG. 5.**
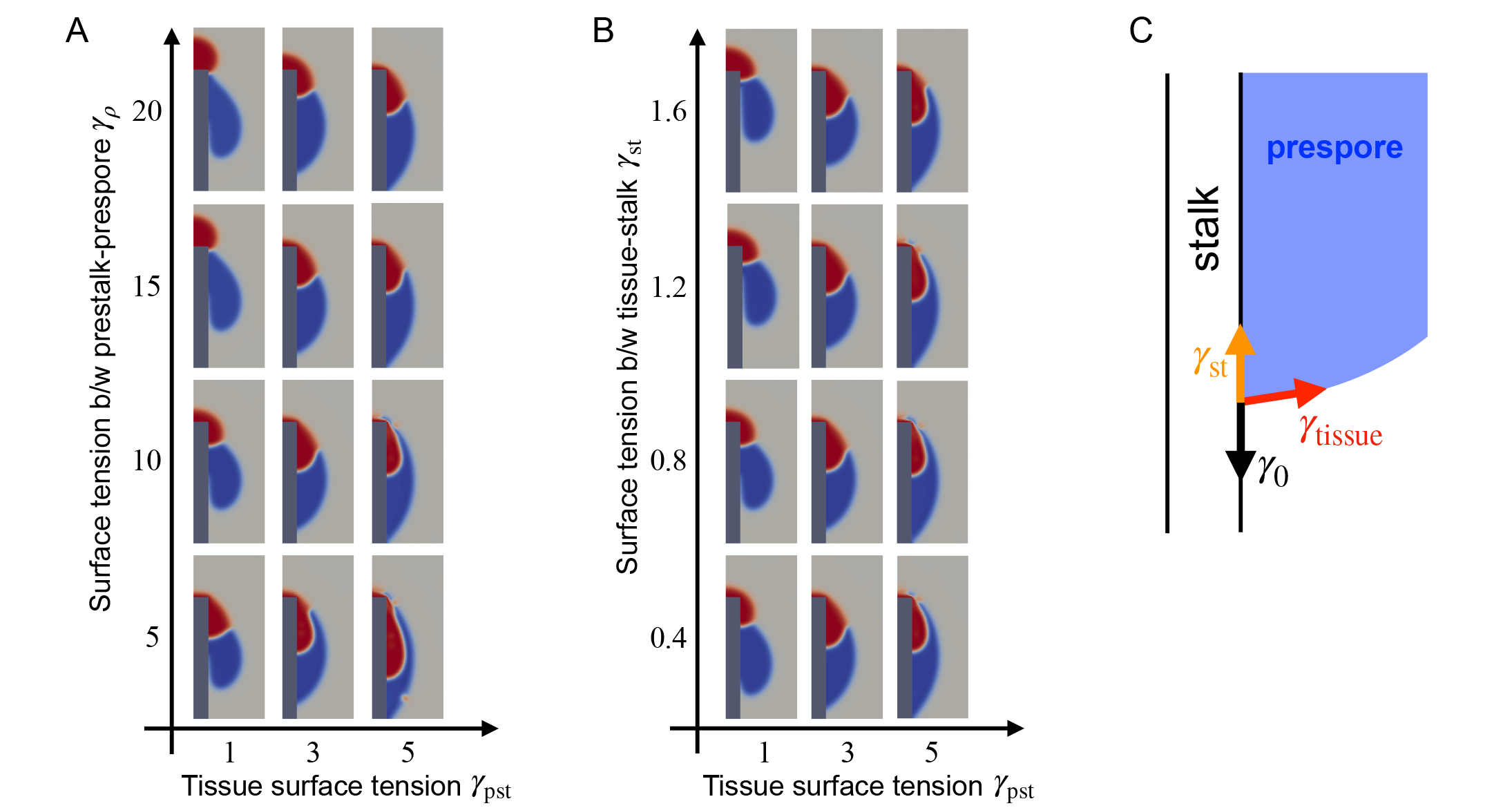
Dependence of tissue morphology on the prestalk–prespore boundary tension *γ*_*ρ*_ and tissue–stalk surface tension *γ*_st_. **A** Simulation results in the *γ*_pst_–*γ*_*ρ*_ plane. **B** Simulation results in the *γ*_pst_–*γ*_st_ plane. Snapshots are taken at *t* = 250 min (A, B). **C** Schematic of the tension balance at the cell mass–stalk contact point.

As mentioned in Sec. III [71], the interfacial tension between prestalk and prespore cells is given by 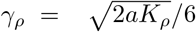. We varied *γ*_*ρ*_ while keeping *K*_*ρ*_*/a* constant so that the prestalk–prespore interface width remained unchanged. Figure 5A summarizes the tissue morphologies obtained in the simulations where *γ*_*ρ*_ and *γ*_pst_ were varied while keeping *γ*_sub_ and *γ*_st_ fixed at *γ*_sub_ = 1.0 and *γ*_st_ = 0.8. The parameter *γ*_*ρ*_ governs how easily the prestalk–prespore boundary can be deformed and thus controls the morphology of this interface. As *γ*_*ρ*_ decreases, the interface becomes longer and more curved. Conversely, increasing *γ*_*ρ*_ makes the interface shorter and straighter, although excessively large *γ*_*ρ*_ relative to *γ*_pst_ leads to separation of the prestalk and prespore regions (Fig. 5A, upper left panel).

Next, we examined the role of the stalk interfacial tension *γ*_st_. Because the values of interfacial tensions between the stalk domain and either the prespore or the prestalk domains are unavailable experimentally, we assigned the same value to both interfaces in the simulations. Figure 5B summarizes the tissue morphologies obtained from simulations in which *γ*_st_ and *γ*_pst_, with *γ*_*ρ*_ = 10 and *γ*_sub_ = 1.0 held constant. The morphology of the posterior end during tissue elevation is governed by the balance between *γ*_st_, which promotes upward deformation of the posterior end, and *γ*_psp_ (= *γ*_pst_*/*2), which resists deformation of the prespore region (Fig. 5B). Consistent with this notion,simulations that successfully exhibit tissue elevation (Fig. 5B; *γ*_pst_ = 1, 3) show that increasing *γ*_st_ increases the height of the posterior end at fixed time points. When *γ*_pst_ is sufficiently large to prevent detach from the substrate, varying *γ*_st_ has no effect on the final morphology (Fig. 5B, fourth column). Collectively, these results demonstrate that both detachment from the substrate and the subsequent cell mass elevation depends on a precise balance of the relevant surface and interfacial tensions.

### F. Effect of disc size on the pre-detachment stage

Our model predicts that mutant strains defective in cell mass detachment should provide insight into the origin of tissue–substrate interfacial tension. In vivo, a thin structure is present between the prespore domain and the substrate that consists of prestalk subtypes that are coated with cellulose and potentially other extracellular matrix proteins [72, 73]. In the *dcsA*-null strain, which lacks cellulose synthase, many culminants failed to complete the early morphogenetic steps, including formation of the basal disc and detachment of the cell mass from the substrate (Fig. 6A; see Methods for details) [69]. Although our model does not explicitly resolve the basal disc or the extracellular matrix, our results suggest that the modeled substrate effectively serves as a mechanical surrogate for these biological interfaces.

**FIG. 6.**
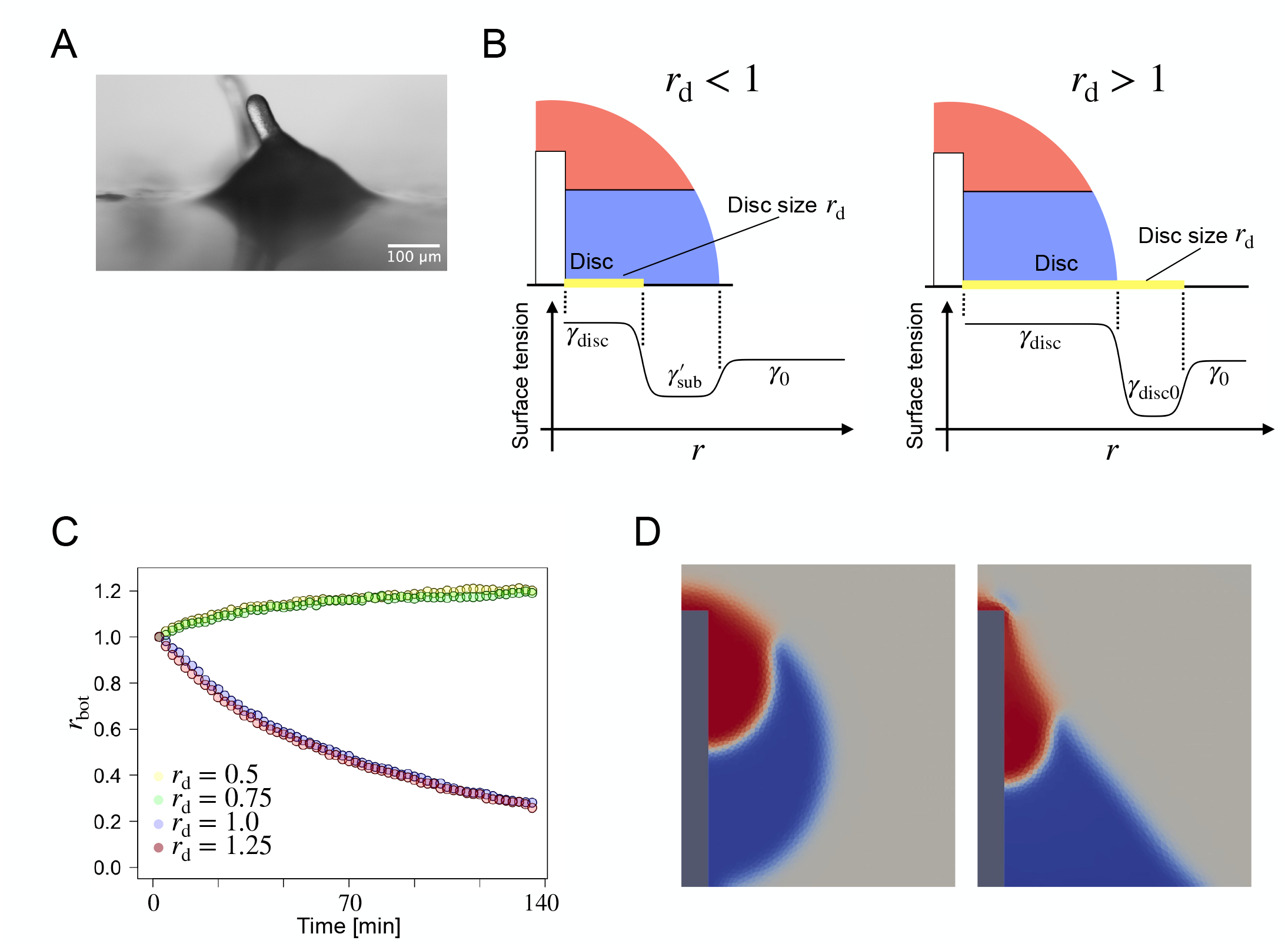
Effect of basal disc on the pre-detachment stage morphology. **A** A representative snapshot of the *dcsA*-null strain. **B** Spatial profile of the effective interfacial tension at the basal tissue–substrate interface, *γ*_bot_(*r*), as a function of *r* in the model. **C** Temporal evolution of the basal contact radius, *r*_bot_(*t*), for different disc sizes *r*_*d*_. **D** Representative morphologies observed in the simulations at *t* = 140 min for disc sizes *r*_*d*_ = 1.25 (left) and *r*_*d*_ = 0.75 (right).

To understand the role of the basal interfaces, we introduce a sub-domain in the substrate and assigned four distinct interfacial tensions (Fig. 6B): *γ*_disc_ for the tissue– basal disc interface, *γ*_disc0_ for the air–basal disc interface, 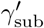 for the tissue–substrate interface, and *γ*_0_ for the air– substrate interface. The effective interfacial tension profile along the basal surface of the tissue and the substrate, *γ*_bot_(*r*), is then given by

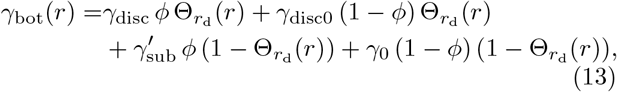

where 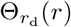 is a step function that equals 1 inside the disc (0 ≤ *r* ≤ *r*_d_) and 0 outside (*r > r*_d_). For the simulations, we set *γ*_disc_ = 2.0, *γ*_disc0_ = 1.2, *γ*_sub_ = 1.2, and *γ*_0_ = 1.6, so that shrinkage of the tissue contact area is favored on the basal disc (*γ*_disc_ *> γ*_disc0_), whereas lateral spreading over the exposed substrate is favored outside the disc 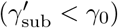. Figure 6C shows the temporal evolution of the basal contact radius, *r*_bot_, for several disc sizes *r*_d_, with all lengths normalized by the initial tissue radius. For large discs (*r*_d_ = 1.0 and 1.25), *r*_bot_ decreased continuously, indicating progressive shrinkage of the tissue–substrate contact region toward detachment (Fig. 6D, left). In contrast, for small discs (*r*_d_ = 0.50 and 0.75), *r*_bot_ initially increased and then approached a plateau, indicating lateral spreading of the basal contact region. Thus, under the interfacial-tension conditions examined here, the transition between spreading and shrinkage occurred between *r*_d_ = 0.75 and 1.0. A snapshot for *r*_d_ = 0.75 showed a morphology that remained attached to the substrate and developed a tent-like profile as stalk elongation proceeded (Fig. 6D, right). Overall, these results indicate that basal-disc extent regulates progression toward cell-mass detachment by controlling the spatial distribution of wettability along the basal interface: a sufficiently large disc favors shrinkage of the basal contact region, whereas exposure to the more wettable substrate favors lateral spreading.

## V. DISCUSSION

Recent studies have revealed that mechanical heterogeneity among tissue regions can organize tissue-scale shape changes, providing a general physical principle in morphogenesis [74, 75]. Spatial differences in surface and interfacial tensions, viscosity, and rigidity have been implicated in zebrafish gastrulation and body-axis elongation [7, 76–78], pollen-tube growth [79, 80], and gonad morphogenesis in *C. elegans* [81]. Here, we extend this principle to *Dictyostelium* culmination, showing that heterogeneous tissue and interfacial mechanics, coupled to stalk-tip extension, are sufficient to generate the characteristic elevation and fruiting-body morphology within a locally force-balanced framework. Our AFM indentation and morphological analyses indicated that surface tension is substantially larger than other mechanical forces acting on the tissue, suggesting that it plays a key role in morphogenesis. Using a mathematical model that explicitly satisfies mechanical force balance, we demonstrated that detachment of the prestalk and prespore cell mass from the substrate and its subsequent elevation can arise spontaneously from the combined effects of surface and interfacial tensions and stalk elongation.

In contrast to previous mathematical models [21– 23, 82], our results suggest that sustained long-range chemotactic migration may be dispensable for upward cell movement during culmination. They further support that cell-mass elevation does not require upward self-propulsion, consistent with earlier replacement experiments showing that mineral oil substituted for spores was also elevated during culmination [37]. This interpretation is also consistent with observations that cAMP waves disappear during culmination [25] and that stage-specific knockdown of ACA, an adenylyl cyclase expressed in the tip region, does not impair culmination [26]. Nevertheless, additional adenylyl cyclases exist besides ACA, and our results do not exclude roles for cAMP signaling in polarization, differentiation, or stalk formation. Rather, they indicate that chemotactic migration need not act as the direct mechanical driver of cell-mass elevation.

Within our model, surface and interfacial tensions, and consequently tissue wettability, play a central role in determining tissue morphology and its dynamics. Successful culmination requires simultaneous mechanical control of three mechanically interfaces: the tissue– substrate interface governing detachment (Fig. 3D), the prestalk–prespore interface maintaining internal organization (Fig. 4D), and the tissue–stalk interface shaping the elevated cell mass (Fig. 5C). Furthermore, our measurements indicate that prestalk region possesses approximately twice the surface tension of prespore region (Sec. II B). Incorporating this experimentally inferred ratio into the simulations reproduces the fruiting body morphology that most closely matches the experimental observations (Sec. IV D). The analysis in Sec. IV F further suggests that basal disc -another prestalk-derived cell population-functionally regulates tissue wettability at the tissue–substrate interface, thereby facilitating cell mass detachment. Collectively, these results indicate that fruiting body morphogenesis is regulated by heterogeneous surface and interfacial tensions, whose spatial organization is established through prestalk cell differentiation. Differential cell motility [83] and adhesion-generated tension gradients [82] have been implicated in formation of the slug and its migration, suggesting that spatially heterogeneous mechanical properties may operate across multiple stages of *Dictyostelium* development.

Our numerical simulations also depicted a variety of aberrant morphogenetic outcomes depending on parameter values. Low tissue surface tension combined with high prestalk–prespore interfacial tension caused complete separation of the two regions (Fig. 5A, *γ*_pst_ = 1.0 and *γ*_*ρ*_ = 15.0 or 20.0), whereas low tissue–substrate interfacial tension combined with high tissue surface tension prevented tissue detachment from the substrate (Fig. 3E, *γ*_pst_ = 5, *γ*_sub_ = 0.5). These results delineate mechanical conditions underlying successful detachment, elevation of the cell mass, and maintenance of prestalk– prespore organization, and provide a framework for interpreting mutant phenotypes. For example, mutations in *trishanku* and *PaxB* cause separation of the prestalk and prespore regions [65–67], qualitatively resembling the separation observed in our simulations. Reducing the modeled basal-disc size impaired shrinkage of the basal contact region and cell-mass detachment but resulted in a tent-like morphology that remained attached to the substrate (Sec. IV F), suggesting a possible mechanical basis for the detachment defect of the *dcsA*-null mutant.

Although our study recapitulates several characteristic features of fruiting body formation in *Dictyostelium*, many questions remain to be addressed. First, several assumptions underlying the present model require further scrutiny. For example, the present model omits other prestalk subtypes, including upper- and lower-cup cells, which play important roles during later stages of fruiting-body morphogenesis. In addition, because stalk formation through prestalk-to-stalk differentiation is a major driver of morphogenesis, a more detailed description of stalk formation will be important. Although the present model represents the stalk as a simple cylinder, tissue morphology can depend on stalk radius (Fig. S3), whereas the actual stalk-tip region has a funnel-like geometry. Future extensions should also incorporate earlier developmental events, including reorientation of the anterior–posterior (prestalk–prespore) axis from the horizontal slug configuration to the vertical early-culminant configuration[52, 84], initial stalk formation from the PstAB core [61], and basal-disc formation [72].

Second, it is unclear how tissue mechanical properties, including surface tension, viscosity, and cellular forces, are regulated at the cellular and molecular levels. In particular, although our results demonstrate that directed migration, such as chemotaxis, is not necessary to reproduce the observed morphogenesis, chemoattractant signaling may still be required for processes such as cell polarization and traction-force generation. Likewise, the mechanical effects of the cellulose-rich extracellular matrix are represented only implicitly through effective surface and interfacial tensions and prescribed boundaries in the present model. Explicitly modeling extracellular-matrix production, deposition, and remodeling will be necessary to connect these tissue-scale mechanical parameters to their cellular and molecular origins [72, 73].

Addressing these issues will require closer integration between theory and experiment. The present modeling framework provides a natural basis for such integration because its predicted velocity field can be quantitatively compared with experimentally measured tissue dynamics. The framework is also extensible, allowing additional cellular processes and active mechanical inputs to be incorporated while retaining a mechanically consistent, locally force-balanced description. Combined with three-dimensional measurements of morphodynamics, such extensions should enable direct connections between cell behaviors, tissue-scale forces, and morphological changes. Ultimately, this approach may provide a mechanistic understanding of how diverse cellular activities are integrated through tissue mechanics to construct the three-dimensional fruiting body and, more broadly, to drive morphogenesis in other systems.

## IV. MATERIAL AND METHODS

### A. Cell culture and development

*Dictyostelium discoideum* wild-type strain Ax4 cells were cultured in HL5 axenic medium supplemented with 1.5% (w/v) glucose, either on Petri dishes or in shaken suspension at 22^°^C. Strains carrying plasmids with neomycin resistance genes were maintained in HL5 medium containing 10 *µ*g/mL G418. Strains carrying a knock-in reporter consisting of PKBR1(N150) fused to tandem mScarlet-I at the *act5* locus were maintained in HL5 medium containing 60 *µ*g/mL hygromycin B. For development, exponentially growing cells were washed twice with phosphate buffer (PB: 12 mM KH_2_PO_4_, 8 mM Na_2_HPO_4_, pH 6.5) and resuspended at a density of 2.0 *×* 10^7^ cells/mL. A 10 *µ*L drop of the cell suspension was placed on a PB agar plate containing 2% purified agar. Alternatively, 5 *µ*L of the suspension was spotted onto a 9 mm^2^ square of membrane filter (A045P047A; ADVANTEC) placed on a piece of filter paper (M-085; ADVANTEC) for stereoscopic imaging during the culmination stage.

### B. Generation of cellulose synthase knockout mutant

The knockout mutant of cellulose synthase was generated in the Ax4 background using CRISPR-Cas9-mediated genome editing. The sgRNA/Cas9 all-in-one vector pTM1285 [85] was obtained from NBRP Nenkin (Japan). sgRNA sequences targeting the 5’ region of the dcsA gene were designed using Cas-Designer [86] and cloned into pTM1285. For construction of the sgRNA expression plasmid, complementary oligonucleotides were synthesized with the following sequences: sense, 5’-AGCACCATCATCTTCAAACTCTCG-3’ ; antisense, 5’-AAACCGAGAGTTTGAAGATGATGG-3’ . The annealed oligonucleotides were inserted into the BpiI site of pTM1285. The resulting targeting plasmid was introduced into Ax4 cells as described previously [87]. Clonal lines were isolated and validated by Sanger sequencing.

### C. Image acquisition

For fluorescence imaging, we followed the protocol described previously [25]. Briefly, an agar block containing developing tissues was excised from the plate and inverted onto a *ϕ*25 mm cover glass (25 mm micro cover glass, Matsunami) mounted on a stainless-steel chamber (Attofluor; Invitrogen). The center of the cover glass was pre-attached with a 50-*µ*m-high ring-shaped spacer (vinyl patch, transparent type Ta-3N, Kokuyo) filled with liquid paraffin (Nacalai Tesque) to match the refractive index. Fluorescence images were acquired using an inverted microscope (IX-81; Olympus) equipped with a multibeam confocal scanning unit (CSU-W1; Yokogawa Electric Corporation, Musashino, Japan) and an EM-CCD camera (iXon Ultra 888; Andor). Optical sections were collected at 1-*µ*m intervals over a total depth of 40 *µ*m using a 20*×* oil-immersion objective lens (UP-LanSAPO 20*×* /0.85 NA; Olympus). Excitation was provided by 561-nm lasers in combination with a dichroic mirror (405/488/561/640) and emission filters (617/73 nm). For stereoscopic imaging, the membrane filter containing developing tissues was transferred onto a PB agar plate prior to time-lapse imaging. An agar block with the membrane filter was then excised and placed on the plate with a 90^°^ rotation to obtain a side view of the culminants. Time-lapse imaging was performed at 1-min intervals using a stereo microscope (SZX16; Evident) equipped with a digital camera (DP23; Evident).

### D. AFM force spectroscopy

AFM force spectroscopy was used to quantify the local mechanical response of the prespore and prestalk regions and the stalk using a NanoWizard 3 Ultra AFM (Bruker) mounted on an Olympus IX70 inverted microscope. Measurements were performed in air at 22°C using OMCL-AC160TS-C3 cantilevers (Olympus; three-sided pyramidal tip, tip radius *<*10 nm; nominal spring constant 26 N/m). The spring constant of the cantilever was calibrated for each experiment using the thermal noise method. Photodetector sensitivity was determined by fitting a linear function to the contact-region slope of a force–distance curve acquired on the glass substrate. On the measurement day, the calibrated deflection sensitivity and spring constant were 29.92 nm/V and 25.15 N/m, respectively, and these values were used to convert cantilever deflection into force. Indentation measurements were performed at an approach and retraction speed of 2 *µ*m/s. Samples were mounted sideways on the substrate to minimize load redistribution caused by bending of the stalk, and at least 16 independent measurements were obtained for each region.

## ACKNOWLEDGMENTS

We acknowledge K. Mitsumoto, K. Sugimura, N. Saito, K. Maki, K. Fuji, and T. Namba for their fruitful discussions and kind support. This work was supported by the Japan Society for the Promotion of Science KAKENHI [Grant Numbers JP22KJ0754 (to S.N.), JP23H04304 (to H.H.), JP23H00384, JP25H01363, JP25H01771, JP25K22490, JP26H00950 (to S.S.), and JP24H01931 (to S.I.)], the Japan Science and Technology Agency CREST [Grant Number JPMJCR1923 (to S.S. and S.I.)] and ACT-X [Grant Number JPMJAX25L8 (to S.N.)].

## DATA AVAILABILITY

The data and simulation code that support the findings of this study are available from the corresponding authors upon reasonable request.

