## Supplementary Information for "Surface Tension and Stalk Elongation Drive *Dictyostelium* Morphogenesis"

### S1 Estimation of tissue Young's modulus and surface tension by AFM

We estimated the Young's modulus $E$ and the surface tension $\gamma$ from force-indentation depth ($F$-$d$) curves measured with atomic force microscopy (AFM). To estimate $E$ and $\gamma$, we performed nonlinear regression using the following relation proposed in prior studies [1,2] (Fig. S1):

$$F = \frac{2}{\pi} E d^2 \tan\phi \left[ 1 + \alpha \left( \frac{s}{d \tan\phi} \right)^{\beta} \right] \tag{S1}$$
$$s = \frac{2\gamma}{E},$$

where $s$ denotes the intrinsic length. The value of $\tan\phi$ is determined by the AFM probe geometry. In this study, we used $\tan\phi = 0.24$. Because previous studies have reported that the coefficients $\alpha$ and $\beta$ are close to unity, we set $\alpha = \beta = 1$ for simplicity. Equation (S1) indicates that the surface tension contribution dominates the response at small indentation depths $d$, whereas the elastic contribution governed by Young's modulus $E$ becomes dominant at larger $d$. To stabilize the nonlinear regression, we first estimated $E$ for indentation depths $d > 50$ nm by setting $\gamma = 0$, and then estimated $\gamma$ for $d < 50$ nm while holding $E$ fixed at the value obtained in the first step.

Using the estimated parameters, we assessed their dependence on sample ID and measurement region by performing a two-way ANOVA with a significance level of 0.05 (Fig. 1E, F). For Young's modulus, significant differences were observed among sample IDs ($p < 0.001$), measurement region ($p < 0.001$), and the interaction effect ($p < 0.001$). For surface tension, only the difference in region was significant ($p = 0.0094$), whereas neither sample ID nor the interaction effect reached significance ($p = 0.052$, $p = 0.78$, respectively).

### S2 Surface-Tension Driven Phase-Field Model

#### *Continuity equations for hydrodynamic phase-field model*

Here we describe the detailed derivation of our phase-field model coupled to a velocity field. As mentioned in the main text, the model consists of three variables: the phase field $\phi(t, \boldsymbol{r})$ representing tissue shape, a variable $\rho(t, \boldsymbol{r})$ representing the fraction of prestalk cells (with the prespore fraction given by $1 - \rho(t, \boldsymbol{r})$, and the velocity field $\boldsymbol{v}(t, \boldsymbol{r})$, where $t$ and $\boldsymbol{r}$

denote time and position, respectively. Briefly, the model is derived by combining the theory of two-component fluid flow [3, 4] with the phase-field method [5].

The volume fractions of prestalk and prespore cells within a tissue are denoted by $\rho\phi$ and $(1-\rho)\phi$, respectively. Let $\boldsymbol{v}_1$ and $\boldsymbol{v}_2$ denote the velocities of prestalk and prespore cells. The time evolution of the volume fractions of the two cell types is described by continuity equations coupled to the flow fields:

$$\frac{\partial(\rho\phi)}{\partial t} + \nabla\cdot(\rho\phi\boldsymbol{v}_1) = 0\ , \tag{S2}$$

$$\frac{\partial(1-\rho)\phi}{\partial t} + \nabla\cdot(1-\rho)\phi\boldsymbol{v}_2 = 0\ . \tag{S3}$$

The mixture (volume-averaged) velocity of the tissue, $\boldsymbol{v}$, and the relative flux $\boldsymbol{j}$ between prespork and prespore cells are defined as

$$\boldsymbol{v} = \rho\boldsymbol{v}_1 + (1-\rho)\boldsymbol{v}_2\ , \tag{S4}$$

$$\boldsymbol{j} = \rho(1-\rho)(\boldsymbol{v}_1 - \boldsymbol{v}_2)\ . \tag{S5}$$

The individual velocities $\boldsymbol{v}_1$ and $\boldsymbol{v}_2$ can be then expressed in terms of $\boldsymbol{v}$ and $\boldsymbol{j}$ as $\boldsymbol{v}_1 = \boldsymbol{v} + \boldsymbol{j}/\rho$ and $\boldsymbol{v}_2 = \boldsymbol{v} - \boldsymbol{j}/(1-\rho)$, and Eqs. (S2) and (S3) lead into

$$\frac{\partial(\rho\phi)}{\partial t} + \nabla\cdot(\rho\phi\boldsymbol{v}) = -\nabla\cdot(\phi\boldsymbol{j}), \tag{S6}$$

$$\frac{\partial(1-\rho)\phi}{\partial t} + \nabla\cdot(1-\rho)\phi\boldsymbol{v} = \nabla\cdot(\phi\boldsymbol{j}). \tag{S7}$$

Notice that Eq. (S6) is identical to Eq. (5) in the main text. Adding Eqs. (S6) and (S7) yields the evolution equation for the tissue phase field $\phi$:

$$\frac{\partial\phi}{\partial t} + \nabla\cdot(\phi\boldsymbol{v}) = 0, \tag{S8}$$

which represents a continuity equation for $\phi$. Under the incompressibility condition $\nabla\cdot\boldsymbol{v} = 0$, this equation is equivalent to the advection form

$$\frac{\partial\phi}{\partial t} + \boldsymbol{v}\cdot\nabla\phi = 0\ . \tag{S9}$$

Since the phase field distinguishes regions inside ($\phi = 1$) and outside ($\phi = 0$) the tissue, this advection equation is consistent with the motion of the deforming tissue boundary moving with velocity $\boldsymbol{v}$.

Instead of directly solving Eq. (S8), we employed a phase-field method in which $\phi$ relaxes toward values of 1 or 0, resulting in distinct domains separated by sharp interfaces [5]. We solved Eq. (2 in the main texts, which is reproduced here for convenience,

$$\frac{\partial\phi}{\partial t} + \nabla\cdot(\boldsymbol{v}\phi) = \Gamma\left(\epsilon\nabla^2\phi - \epsilon^{-1}G'(\phi) + c\epsilon|\nabla\phi|\right)\ . \tag{S10}$$

$G_\phi(\phi) = \phi^2(\phi-1)^2$. As mentioned in the main text, $\phi$ is the phase field for the tissue domain ($\phi = 1$ inside and $\phi = 0$ outside). Equation (S10) describes advection by $\boldsymbol{v}$, relaxation toward the minima of $G_\phi$, and the term $c\epsilon|\nabla\phi|$ cancels the curvature contribution in $\nabla^2\phi$, suppressing the intrinsic curvature-driven motion of the phase-field interface. Since the right-hand side of the above equation relaxes to zero to a good approximation once the system separates into $\phi = 0$ and $\phi = 1$ domains, this equation is effectively equivalent to Eq. (S8). In numerical simulation, we employed further improved method [6] to solve the phase-filed equation (see Sec. S3).

Finally, the evolution equation for $\rho$ is obtained from Eqs. (S8) and (S6) as

$$\frac{\partial\rho}{\partial t} + \boldsymbol{v}\cdot\nabla\rho = -\frac{1}{\phi}\nabla\cdot(\phi\boldsymbol{j})\ . \tag{S11}$$

***Stress and chemical potential***

The force $\boldsymbol{f}_a$ and the relative flux $\boldsymbol{j}$ (see Eqs. (3), (5), (8), and (9) in the main text) are derived from the free energy given in Eq. (6) of the main text by invoking the energy dissipation law $dF/dt \leq 0$. The time derivative of the free energy is expressed as a sum of force–flux products,

$$\frac{dF}{dt} = \int \{\nabla \boldsymbol{v} : \sigma + \phi \boldsymbol{j} \cdot \nabla \mu\}\, dV. \tag{S12}$$

where $\nabla \boldsymbol{v}$ is the strain rate tensor.

The quantities $\sigma$ and $\mu$ are interpreted as the stress tensor and the chemical potential difference between prestalk and prespore cells (up to a factor of $\phi$; $\mu$ is refereed to as chemical potential below), respectively, both derived from the free energy. Explicit expressions for $\sigma$ and $\mu$ are derived below.

Our free energy $F$ is given by $F = F_\rho + F_1 + F_2 + F_3$ (Eq. (6) in the main text), where

$$F_\rho = \int \{\phi g(\rho, \nabla\rho) + (1-\phi) f_\rho\}\ , \tag{S13}$$

$$F_1 = \int_{\Gamma_{\mathrm{tissue}}} \gamma_{\mathrm{tissue}}(\rho) dS = \int \gamma_{\mathrm{tissue}} |\nabla \phi|\, dV\ , \tag{S14}$$

$$F_2 = \int_{\Gamma_{\mathrm{sub}}} \gamma_{\mathrm{sub}} dS\ , \tag{S15}$$

$$F_3 = \int_{\Gamma_{\mathrm{st}}} \gamma_{\mathrm{st}} dS\ , \tag{S16}$$

and

$$g(\rho, \nabla\rho) = G_\rho(\rho) + \frac{K_\rho}{2} |\nabla \rho|^2\ . \tag{S17}$$

$$\gamma_{\mathrm{tissue}}(\rho) = \rho\gamma_{\mathrm{pst}} + (1-\rho)\gamma_{\mathrm{psp}} \tag{S18}$$

First, we consider the time evolution of $F_\rho$:

$$\begin{aligned}
\frac{dF_\rho}{dt} &= \int \frac{\partial}{\partial t}\left[\phi g + (1-\phi) f_\rho\right] dV \\
&= \int \left\{ (g - f_\rho)\partial_t \phi + \phi \left( \frac{\partial g}{\partial \rho} \partial_t \rho + \frac{\partial g}{\partial(\partial_k \rho)} \partial_t \partial_k \rho \right) \right\} dV \\
&= \int \left\{ -(g - f_\rho) v_k \partial_k \phi - \left( \phi \frac{\partial g}{\partial \rho} - \partial_i \left( \phi \frac{\partial g}{\partial(\partial_i \rho)} \right) \right) \left( v_k \partial_k \rho + \frac{1}{\phi} \nabla \cdot (\phi \boldsymbol{j}) \right) \right\} dV \\
&= \int \left\{ \left( (\phi g + (1-\phi) f_\rho)\delta_{ik} - \phi \frac{\partial g}{\partial(\partial_i \rho)} \partial_k \rho \right) \partial_i v_k + \phi j_k \partial_k \left( \frac{\partial g}{\partial \rho} - \phi^{-1} \partial_i \left( \phi \frac{\partial g}{\partial(\partial_i \rho)} \right) \right) \right\} dV.
\end{aligned}$$

Here we have used Eqs. (S11) and (S8), together with integration by parts. Partial derivatives with respect to time and space are denoted by $\partial_t$ and $\partial_k$, respectively, where the index $k$ labels spatial coordinates, and the Einstein summation convention is implied for repeated indices. Therefore stress and chemical potential caused by $F_\rho$ are determined as

$$\sigma_\rho = (\phi g + (1-\phi) f_\rho)\boldsymbol{I} - K_\rho \phi \nabla\rho \otimes \nabla\rho \tag{S19}$$

$$\mu_\rho = \frac{\partial g}{\partial \rho} - \phi^{-1} \partial_i \left( \phi \frac{\partial g}{\partial(\partial_i \rho)} \right) = \frac{\partial g}{\partial \rho} - K_\rho \nabla^2 \rho - K_\rho \phi^{-1} \nabla\phi \cdot \nabla\rho \tag{S20}$$

Note that isotropic part of the stress (component proportional to $\boldsymbol{i}$) has no physical consequence under the incompressibility condition, $\nabla \cdot \boldsymbol{v}$, and we can omit the first term of $\sigma_\rho$. The second term is non-zero at the prespore-prestalk interface, indicating the interfacial stress.

Next, we calculate the time derivative $dF_1/dt$ (Eq. (S14)), originating from the tissue surface free energy $\gamma_{\rm tissue}$. The result is

$$\frac{dF_1}{dt} = \int \left\{ \gamma_{\rm tissue} \left( |\nabla\phi|\delta_{ik} - \frac{\partial_i\phi\partial_k\phi}{|\nabla\phi|} \right) \partial_i v_k + \phi j_k \partial_k \left( \phi^{-1} \frac{\partial\gamma_{\rm tissue}}{\partial\rho} |\nabla\phi| \right) \right\} dV \ ,$$

and thus

$$\sigma_1 = \gamma_{\rm tissue} \left( |\nabla\phi| \boldsymbol{I} - \frac{\nabla\phi\otimes\nabla\phi}{|\nabla\phi|} \right) \quad \text{(S21)}$$

$$\mu_1 = \phi^{-1}\frac{\partial\gamma_{\rm tissue}}{\partial\rho}|\nabla\phi| \quad \text{(S22)}$$

Finally, we derive the expressions for the stresses arising from $\gamma_{\rm sub}(\phi)$ and $\gamma_{\rm st}(\phi)$. Both surface tensions are defined as $\phi\,\gamma_{\rm s} + (1-\phi)\,\gamma_0$, where $\gamma_{\rm s}$ denotes the interface-specific surface tension ($\gamma_{\rm s} = \gamma_{\rm sub}$ on $\Gamma_{\rm sub}$ and $\gamma_{\rm s} = \gamma_{\rm st}$ on $\Gamma_{\rm st}$), while $\gamma_0$ denotes the surface tension between the substrate or stalk and air. As mentioned in the main text, we assume that the surface tension between the tissue and the stalk is identical for prestalk and prespore cells. The corresponding stress tensors are expressed as

$$\sigma_2 = \gamma_{\rm sub}\left(\boldsymbol{I} - \boldsymbol{n}_{\rm sub}\otimes\boldsymbol{n}_{\rm sub}\right)\delta_{\Gamma_{\rm sub}}(\boldsymbol{r}) \ , \quad \text{(S23)}$$

$$\sigma_3 = \gamma_{\rm st}\left(\boldsymbol{I} - \boldsymbol{n}_{\rm st}\otimes\boldsymbol{n}_{\rm st}\right)\delta_{\Gamma_{\rm st}}(\boldsymbol{r}) \ . \quad \text{(S24)}$$

Here, $\boldsymbol{n}_{\rm sub}$ and $\boldsymbol{n}_{\rm st}$ denote the unit normal vectors to the substrate and stalk interfaces, respectively, and $\delta_{\Gamma_{\rm sub}}$ and $\delta_{\Gamma_{\rm st}}$ are surface delta functions supported on the corresponding interfaces. These surface tensions do not give rise to any chemical potential.

Collecting stresses and chemical potentials derived above, we obtain

$$\sigma = -K_\rho\phi\nabla\rho\otimes\nabla\rho + \gamma_{\rm tissue}|\nabla\phi|\left(\boldsymbol{I} - \boldsymbol{n}\otimes\boldsymbol{n}\right) + \sigma_2 + \sigma_3 \ , \quad \text{(S25)}$$

$$\mu = \frac{\partial g}{\partial\rho} - K_\rho\nabla^2\rho - K_\rho\phi^{-1}\nabla\phi\cdot\nabla\rho + \phi^{-1}\frac{\partial\gamma_{\rm tissue}}{\partial\rho}|\nabla\phi| \ . \quad \text{(S26)}$$

where we can omit the isotropic component of stress.

#### *Derivation of $\boldsymbol{f}_a$ and $\boldsymbol{j}$*

We determine the force $\boldsymbol{f}_a$ and the relative flux $\boldsymbol{j}$ within the linear-response regime so as to ensure energy dissipation $dF/dt \leq 0$ [4, 7] and to satisfy appropriate boundary conditions. To determine the force $\boldsymbol{f}_a$, the stress $\sigma$ is incorporated into the Stokes equation as

$$\boldsymbol{f}_a = -\nabla\cdot\boldsymbol{\sigma}, \quad \text{(S27)}$$

by which the first term in Eq. (S12) contributes negatively to $dF/dt$, ensuring dissipation. By substituiting Eq. (S25) into Eq. (S27), we obtain the explicit form of $\boldsymbol{f}_a$ as

$$\begin{aligned}\boldsymbol{f}_a &= \nabla\cdot\left[K_\rho\phi\nabla\rho\otimes\nabla\rho - \gamma_{\rm tissue}|\nabla\phi|\left(\boldsymbol{I} - \boldsymbol{n}\otimes\boldsymbol{n}\right) + \sigma_2 + \sigma_3\right] \\ &= \nabla\cdot\left(K_\rho\phi\nabla\rho\otimes\nabla\rho\right)|\nabla\phi|\left(\gamma_{\rm tissue}c\boldsymbol{n} - (\boldsymbol{I} - \boldsymbol{n}\otimes\boldsymbol{n})\nabla\gamma_{\rm tissue}\right) - \nabla\cdot\left(\sigma_2 + \sigma_3\right) \quad \text{(S28)}\end{aligned}$$

that is used in the numrical simulations (Eq. (8) in the main text).

For the relative flux $\boldsymbol{j}$, we adopted

$$\boldsymbol{j} = -L(\phi,\rho)\,\nabla\mu \quad \text{(S29)}$$

with positive kinetic coefficient $L(\phi,\rho)$, which ensures that the second term in Eq. (S12) contributes negatively to $dF/dt$. This expression simply reflects that the relative flux occurs along the gradient of the chemical potential. The kinematic coefficient $L(\phi,\rho)$ is chosen to satisfy $\boldsymbol{j} \propto \rho(1-\rho)$ to be compatible with the definition Eq. (S5). In addition, both $\boldsymbol{v}_1$ and $\boldsymbol{v}_2$, the velocities of prestalk and prespore cells, must be parallel to the tissue surface boundary to avoid leakage of cells to outside of the tissue. We take $L(\phi,\rho) = \alpha_{\rm j}\rho(1-\rho)\left(I - \boldsymbol{n}\otimes\boldsymbol{n}\right)$ with a positive constant $\alpha_{\rm j}$ and $\boldsymbol{n} = -\nabla\phi/|\nabla\phi|$. We finally obtain the flux

$$\boldsymbol{j} = -\alpha_{\rm j}\rho(1-\rho)\left(I - \boldsymbol{n}\otimes\boldsymbol{n}\right)\nabla\left(\frac{\partial g}{\partial\rho} - K_\rho\nabla^2\rho - K_\rho\phi^{-1}\nabla\phi\cdot\nabla\rho + \phi^{-1}\frac{\partial\gamma_{\rm tissue}}{\partial\rho}|\nabla\phi|\right) \quad \text{(S30)}$$

that is used in the numerical simulations (Eq. (9) in the main text).

### S3 Practical simulation method to solve the phase-field equation

In practice, to perform stable and accurate numerical integration of the phase-field equation, Eq. (S10), we employed the numerical scheme proposed by Badillo et al. [6]. This method splits the phase-field dynamics at each time step into two subproblems: first, a smoothing (or reinitialization) equation for the phase-field is solved, followed by the solution of a transport equation.

The first subproblem consists of solving the following equation:

$$\frac{\partial \phi}{\partial \tau} = \nabla \cdot \left\{ \epsilon^2 \nabla \phi - \epsilon \sqrt{2}\, \phi(1-\phi) \frac{\nabla \phi}{|\nabla \phi|} \right\}, \tag{S31}$$

where $\tau$ denotes a computational pseudo-time that does not correspond to physical time. It has been shown that the stationary profile of $\phi$ obtained by solving this equation coincides with that obtained from Eq. (S10) in the absence of the advection term [6]. Within each update of $\phi$, we integrate this equation in pseudo-time $\tau$ until $d\phi/d\tau$ becomes sufficiently small, while keeping the other variables fixed. This procedure yields a phase-field profile with a sufficiently sharp interface, corresponding to a well-defined tissue boundary.

The second subproblem is the transport and deformation of $\phi$, for which we solve the advection equation

$$\frac{\partial \phi}{\partial t} + \boldsymbol{v} \cdot \nabla \phi = 0 \ . \tag{S32}$$

Note that this equation is identical to the continuity equation Eq. (S8) under the incompressibility condition. Both Eqs. (S31) and (S32) preserve the total cell mass, $\int_\Omega \phi \, dV$, except outflow of prestalk cell by differentiation at the the stalk tip.

### S4 Stalk radius and tissue morphology

In the numerical simulations in the main text we fixed the stalk radius as $r_{\mathrm{st}} = 3$. To test whether the stalk radius affects tissue morphology, we performed numerical simulations varying the stalk radius $r_{\mathrm{st}}$ along with the prestalk viscosity $\eta_{\mathrm{pst}}$. We set the relative surface tension to $\gamma_{\mathrm{pst}}/\gamma_{\mathrm{psp}} = 1.05$, and all other parameters were identical to those used in Sec. 4.4 of the main text. As shown in Fig. S3, the stalk radius $r_{\mathrm{st}}$ controls the width of the tissue tip morphology. When the stalk is thin, the tissue is stretched along the stalk and a more pronounced protruding tip forms (first column of Fig. S3). As the prestalk viscosity $\eta_{\mathrm{pst}}$ increases, the tissue narrows more abruptly in the necking region. These results suggests that tissue morphology is determined not only by the viscosity and surface tension contrasts between the prestalk and prespore domains, but also by the stalk radius.

Table S1: Geometric measurements of fruiting bodies ($n = 4$). Values in the final row are the mean ± SD.

| Sample ID | Stalk length $L$ ($\mu$m) | Cell-mass radius $R$ ($\mu$m) | Stalk radius $r$ ($\mu$m) |
|---|---|---|---|
| Sample 1 | 539 | 89 | 8.2 |
| Sample 2 | 808 | 84 | 10.3 |
| Sample 3 | 679 | 95 | 8.1 |
| Sample 4 | 793 | 123 | 16.2 |
| Mean ± SD | $705 \pm 125$ | $98 \pm 17$ | $10.7 \pm 3.8$ |

Table S2: Parameters used in the present study.

| Symbol | Meaning | Value / explored range |
|---|---|---|
| *Physical parameters used in force estimates* | | |
| $r$ | Stalk radius | $1.07 \times 10^{-5}$ m |
| $L$ | Stalk length | $7.1 \times 10^{-4}$ m |
| $R$ | Radius of the cell mass | $1.0 \times 10^{-4}$ m |
| $\rho_{\mathrm{m}}$ | Cell mass density used in the gravitational-force estimate | $1.0 \times 10^{3}\,\mathrm{kg\,m^{-3}}$ |
| $E$ | Young's modulus of the stalk | $8.0 \times 10^{7}$ Pa |
| $g$ | Gravitational acceleration | $9.8\,\mathrm{m\,s^{-2}}$ |
| *Continuum-model parameters* | | |
| $r_{\mathrm{st}}$ | Stalk radius in simulations | 1–5 |
| $\Gamma$ | Coefficient in the phase-field equation for $\phi$ | 1/6 |
| $\epsilon$ | Interface-width parameter in the phase-field description | 5/6 |
| $a$ | Double-well coefficient in $G_\rho(\rho) = a\rho^2(1-\rho)^2$ | Varied with $\gamma_\rho$ while keeping $K_\rho/a$ fixed |
| $K_\rho$ | Gradient-energy coefficient for the cell-type field $\rho$ | Varied proportionally with $a$ while keeping $K_\rho/a$ fixed |
| $\gamma_{\mathrm{pst}}$ | Surface tension of the prestalk region | 1.0–7.0 |
| $\gamma_{\mathrm{psp}}$ | Surface tension of the prespore region | $0.5\gamma_{\mathrm{pst}}$, $0.95\,\gamma_{\mathrm{pst}}$ |
| $\gamma_{\mathrm{tissue}}(\rho)$ | Local tissue surface tension | $\rho\gamma_{\mathrm{pst}} + (1-\rho)\gamma_{\mathrm{psp}}$ |
| $\gamma_\rho$ | Interfacial tension between prestalk and prespore regions | 5–20 |
| $\gamma_{\mathrm{sub}}$ | Tissue–substrate interfacial tension | 0.5–2.0 |
| $\gamma_{\mathrm{st}}$ | Tissue–stalk interfacial tension | 0.4–1.6 |
| $\alpha_{\mathrm{j}}$ | Mobility coefficient for the relative prestalk–prespore flux | 1 |
| $v^{\mathrm{in}}$ | Prestalk-cell influx speed relative to the moving stalk-tip boundary | $2 \times 10^{-5}$ |
| $\dot{z}^{\mathrm{top}}$ | Upward velocity of the stalk-tip boundary | $r_{\mathrm{ve}}v^{\mathrm{in}} = 5.0 \times 10^{-5}$ |
| $r_{\mathrm{ve}}$ | Volumetric expansion ratio upon prestalk-to-stalk differentiation and vacuolation | 2.5 |
| $\eta_v$ | Viscosity of the tissue | $1.0 \times 10^{3}$ |
| $\eta_{\mathrm{pst}}/\eta_{\mathrm{psp}}$ | Viscosity ratio (prestalk region / prespore region) | 1–99 |
| $\eta_{\mathrm{ext}}$ | Viscosity of the exterior phase | $\eta_{\mathrm{psp}}$ |
| $r_{\mathrm{d}}$ | Basal-disc radius normalized by the initial tissue radius | 0.5–1.25 |
| $\gamma_{\mathrm{disc}}$ | Tissue–basal-disc interfacial tension | 2.0 |
| $\gamma_{\mathrm{disc0}}$ | Air–basal-disc interfacial-energy referencen | 1.0 |
| $\gamma'_{\mathrm{sub}}$ | Tissue–substrate interfacial tension outside the basal disc | 1.2 |
| $\gamma_0$ | Air–substrate interfacial-energy reference | 1.6 |
| *Numerical settings* | | |
| $L_r$ | Radial size of the computational domain | 50 |
| $L_z$ | Axial size of the computational domain | 150 |
| $\Delta t$ | Numerical time step | 0.1 |

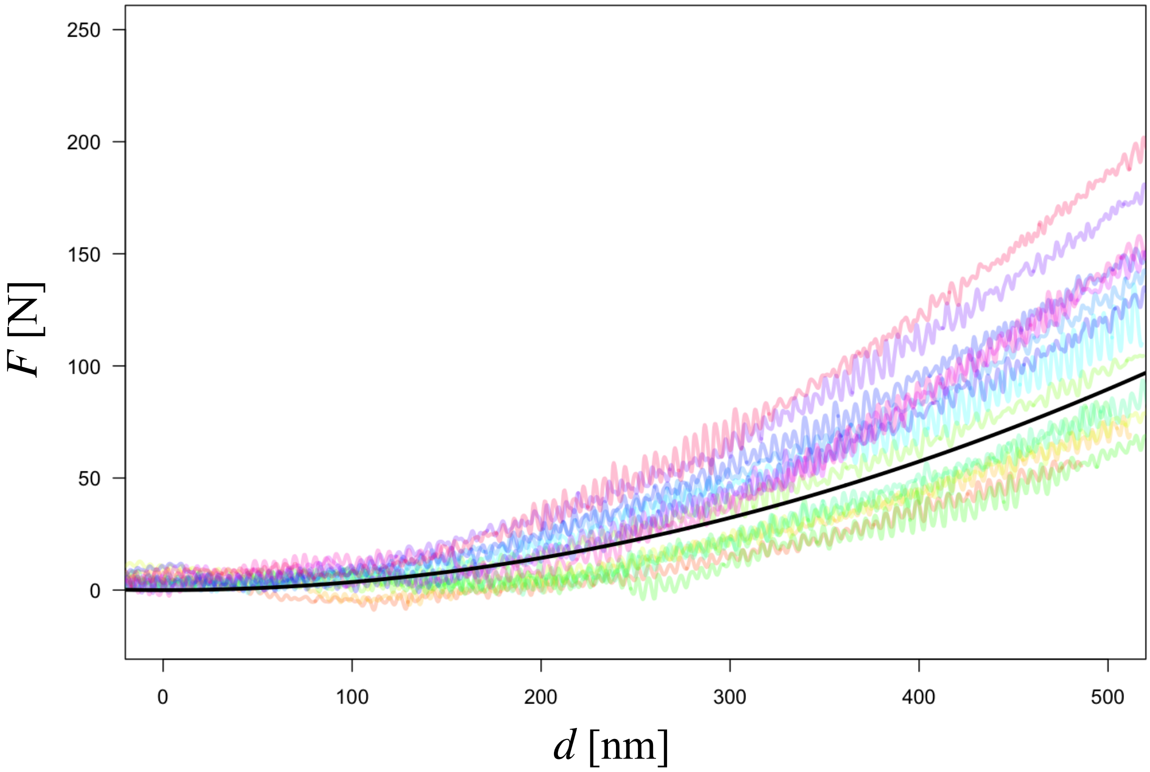


Figure S1: A typical force versus indentation curve. The black curve shows the theoretical relation given by Eq. (S1). Colored curves show repeated measurements collected from the same tissue, with each color corresponding to one measurement repeat.

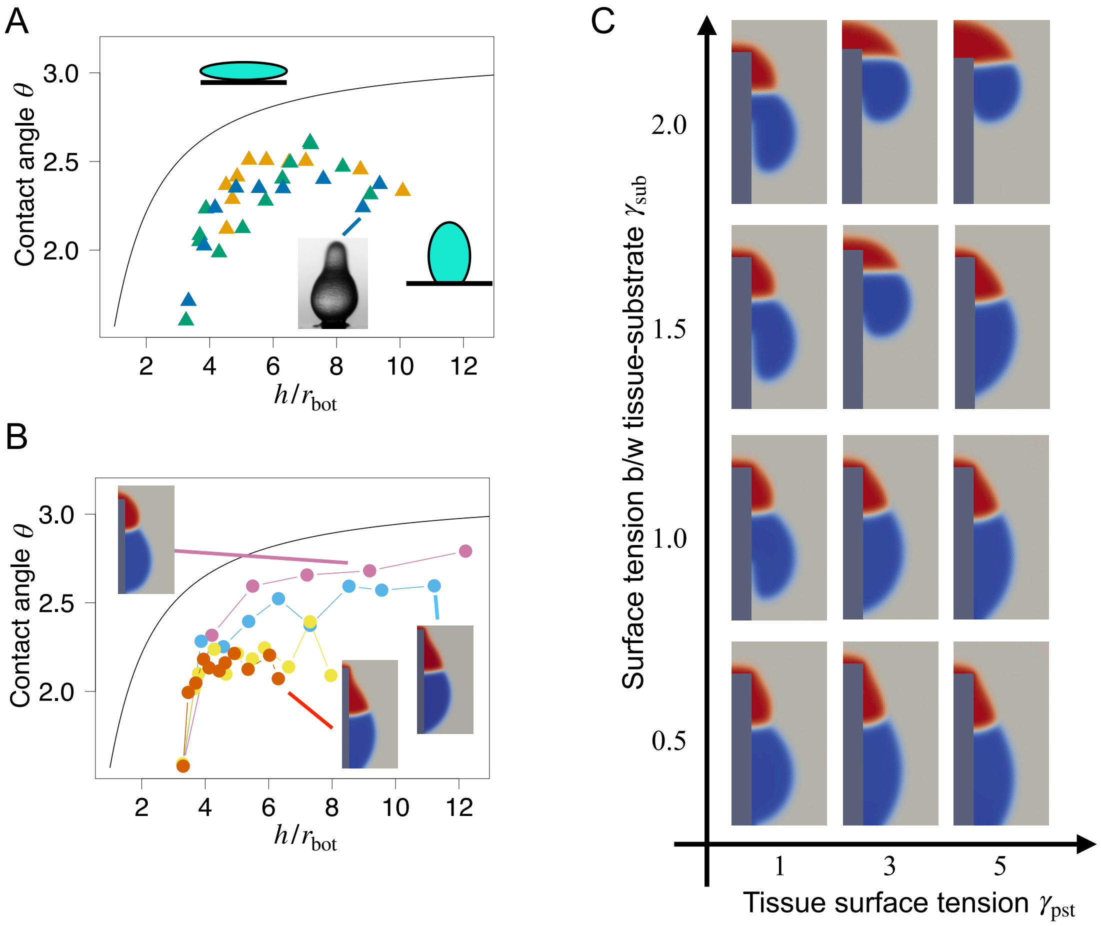


Figure S2: Comparison of experimental and simulation results in early developmental stage with $\gamma_{\text{psp}} = 0.95\,\gamma_{\text{pst}}$. **A**, **B** Plots of the base radius to height ratio $h/r_{\text{bot}}$ versus the contact angle $\theta$ in experimental observations and simulations. The spherical cap curve is shown in black for reference; the tissue shape is more prolate above the curve and more oblate below it. **A** is the same as that shown in Fig. 3B in the main text. In **B**, colors indicate the value of $\gamma_{\text{pst}}$, with red, yellow, light blue, and pink corresponding to 7.0, 5.0, 3.0, and 1.0, respectively. **C** Phase diagram from simulations on the ($\gamma_{\text{pst}}$-$\gamma_{\text{sub}}$) plane with $\gamma_{\text{psp}} = 0.95\,\gamma_{\text{pst}}$.

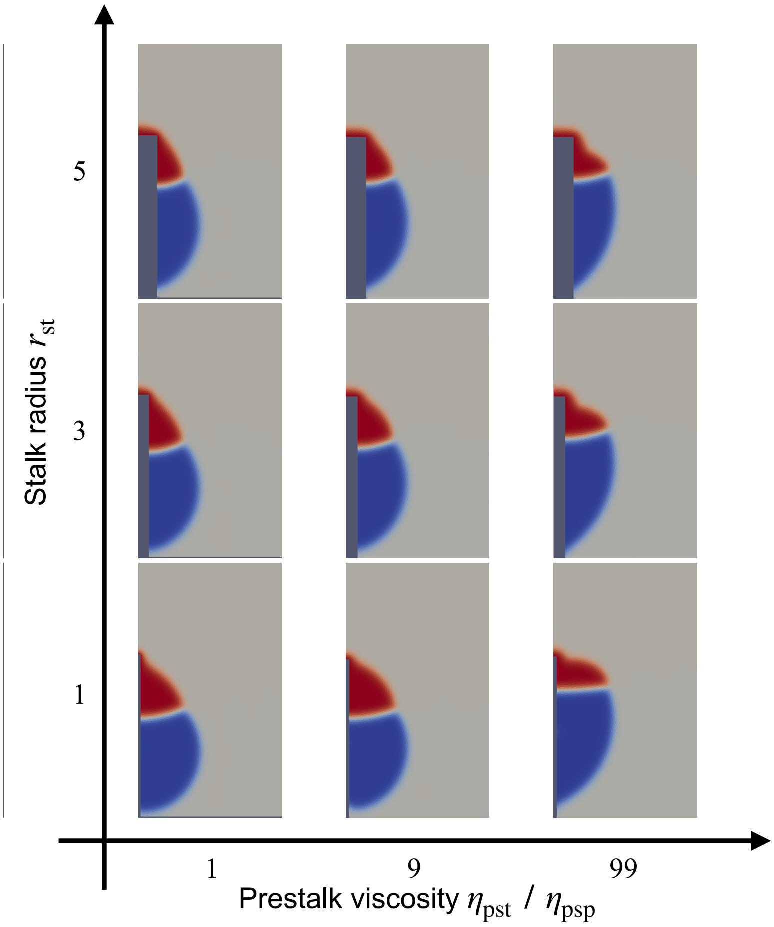


Figure S3: Viscosity and stark radius govern tissue morphology. Phase diagram from simulations on the $(\eta_{pst}/\eta_{psp})$–$(\gamma_{pst}/\gamma_{psp})$ plane.
